# Genomic signals of local adaptation in UK black alder populations (*Alnus glutinosa*)

**DOI:** 10.64898/2026.10.15.759341

**Authors:** Phoebe Swift, Mike Charters, Rômulo Carleial, Patricia Wood, Louise Gathercole, Ted Chapman, Richard Buggs

## Abstract

Black alder (*Alnus glutinosa*) is a native tree species in the British Isles, which it colonised after the last glaciation from continental refugia. It is now a widespread pioneer species, thriving on wetlands and riverbanks. We present whole genome sequence data for 324 alder seedlings grown from seed collected from 44 populations across the UK. Among these we found two triploids and one sample with trisomy at chromosome five. Removing these, we found little population genetic structure apart from diverging northern populations with lower diversity. Using Latent-Factor Mixed Models (LFMM) analyses we identified 1,364 loci whose allele frequencies were significantly associated with environmental variables. Of these loci, 93% were associated with the number of snow days. In contrast, the greatest total genomic signal (variance in SNP effect sizes of 7.38 million SNP loci) was observed for mean temperature of the wettest quarter and annual number of frost days. Estimates of maladaptation using 1.49 million SNPs, predicted that northern populations are worst adapted to present and future worst-case climates. Our results suggest that similarity of source-site temperature variables should be prioritised when choosing seed sources for new planting, and northern populations could benefit from assisted gene flow from the south.

## Introduction

It is widely assumed that a population persisting for many generations in an environment will become adapted to local conditions due to local selection pressures (Kawecki and Ebert 2004; Whitlock 2015). This adaptation arises from standing genetic variation and *de novo* mutations (Savolainen et al. 2013). Genome-environment association (GEA) studies, which compare SNP allele frequencies between populations of the same species along environmental gradients, aim to identify loci involved in local adaptation (Lasky et al. 2023; Rellstab et al. 2015). This knowledge can be used to predict maladaptation of populations under current and future climates, and inform restoration programs aimed at ‘rescuing’ vulnerable populations through assisted gene flow or migration (Beheregaray et al. 2026). Modern high-throughput sequencing technologies have enabled fast and cheap whole-genome sequencing (WGS) of hundreds, sometimes thousands, of individuals, making genome-wide GEA studies possible (Long et al. 2025; Sang et al. 2022; Liang et al. 2025; Lazic et al. 2024).

The UK has ambitious tree planting targets to meet its Net Zero commitments (Committee on Climate Change 2019), and also improve biodiversity and flood management. Success relies on trees surviving at their planting sites in present and future climates. Rapid assessment of planting stock suitability can be achieved by UK-scale GEA studies, to characterise current local adaptation and identify seed sources adapted to present and future climates at planting sites.

Black alder trees (*Alnus glutinosa*) are monoecious, wind-pollinated, pillars of ecosystems scattered across Europe (McVean 1953; De Kort et al. 2014). They are a riparian pioneer species, commonly used to restore degraded marsh and wetland landscapes, increasing biodiversity (Gomes Marques et al. 2022; Claessens et al. 2010; Vaht et al. 2010). Their root networks are vital in building up river banks, preventing erosion, and forming symbiotic relationships with nitrogen-fixing bacteria which fertilises the soil for ecosystem establishment (Forget et al. 2013; Ferguson and Bond 1953; Huang et al. 2026). This capacity to stabilise riverways makes them important in flood defence and afforestation (Vacek et al. 2022). Mature alder are stress-tolerant to drought and heat, and are effective at absorbing atmospheric pollutants such as nitrogen oxides, making them good candidates for planting in urban settings (Zhang et al. 2020; Vaht et al. 2010).

Riparian ecosystems, which depended on water and precipitation to function, are particularly sensitive to flooding, droughts and temperature extremes associated with climate change (Nakamura 2022). Species in these habitats must adapt quickly to cope with prolonged periods of stress. Alder trees are also threatened by fungal diseases such as root rot caused by *Phytophthora* species (Huang et al., 2026), and habitat loss due to agriculturally-driven land-use change (Beatty et al. 2015).

Despite roles in flood management and habitat restoration, research on UK alder populations’ local adaptation and potential future response to climate change is lacking, in contrast to mainland Europe: (Cox et al. 2011; De Kort et al. 2014; Vacek et al. 2022). Not being a major timber-producing species, alder does not receive the same level of funding and interest as other tree species such as oak and birch. This has created knowledge gaps, which may become detrimental if afforestation with alder continues without assessment of its resilience against future threats.

Analysis of the genetic structure of *A. glutinosa* in Europe using chloroplast sequence data and nuclear microsatellites (Havrdová, et al., 2015) suggests British populations are similar to those in France, Germany and the Low Countries, and expanded from refugia in the east of the Iberian Peninsula after the last glaciation (Vít et al. 2017). Population structure and GEA studies on alder in Europe primarily used nuclear and/or chloroplast microsatellites, amplified fragment length polymorphisms, or reduced-representation sequencing approaches such as ddRAD or genotyping-by-sequencing (Beatty et al. 2015; Cox et al. 2011; Hrivnák et al. 2023; De Kort et al. 2014). These studies demonstrate high within-population genetic diversity and low genetic differentiation among populations, likely due to wind-dispersed pollen and water-dispersed seeds enabling long-distance gene flow. Despite this connectivity, genome scans and GEAs in Europe identified signatures of local adaptation along climate gradients, with SNPs associated with hydrological regimes and temperature variables (Cox et al. 2011; Hrivnák et al. 2023; De Kort et al. 2014). De Kort et al. (2014) combined GEA and common garden experiments to show that temperature-associated SNPs also covaried with variation in leaf size and phenology, providing additional evidence that these loci contribute to local adaptation.

The alder populations used in this study were collected as part of the UK National Tree Seed Project (UKNTSP), which aims to preserve the natural genetic variation of UK native tree species by collecting and storing seeds from across the native range (Chapman 2024). These collections represent an important scientific resource, enabling the study of local adaptation (through genetic analyses and germination tests), the characterisation of genetic diversity, and the provision of material for provenance and breeding trials. Through whole genome re-sequencing of seedlings grown from UKNTSP collections, we identify populations within the UKNTSP collections containing polyploids and trisomy. We investigate variables driving genetic variation, and uncover clines in allele frequency between populations and environmental extremes to identify loci with the potential to enhance adaptation through assisted gene flow protocols, or targeted planting (matching genotype with environment).

## Material and Methods

### Plant material, DNA Extraction and Sequencing

Alder seeds were collected between 2014 and 2019 as part of the United Kingdom National Tree Seed Project (UKNTSP). Seeds were dried at 15% relative humidity before being stored at -20°C in the Millenium Seed Bank (MSB), Surrey. In November 2021, seeds from 44 populations containing 1-26 mother trees each (mean 6.8 ± 5.5 SD, Table S1) were withdrawn and transported to the Royal Botanic Gardens, Kew.

Seeds were sown onto a damp 3:1 growing medium of Kew Pot (25% Coir, 35% Composted Bark, 15% Sterilised Soil, 5% Grit, 10% Sand, 10% Wood Fibre) and perlite. Germination occurred in the Kew Jodrell glasshouse under a 12-hour light-dark cycle with 20°C ambient temperature and LED lights (350 W/m2) during the day, and 17°C at night. Relative humidity was maintained below 60% through irrigation and ventilation. Seedlings were fertilised with liquid feed and harvested after 10 weeks using sterilised forceps.

DNA extractions were conducted by Edinburgh Genetics Limited using their modified CTAB protocol (personal correspondence). One seedling was selected per mother tree. Up to 150mg of leaf tissue was frozen in liquid nitrogen, ground into a powder, and incubated in 1ml of CTAB lysis buffer at 65°C for 1hr. An equal volume of chloroform/isoamyl alcohol (24:1) was added, and the DNA was precipitated using isopropyl alcohol and 3M sodium acetate via incubating at -20°C for 2hrs. The DNA pellet was washed twice with 75% ethanol and resuspended in up to 100µL of TE buffer. DNA quality was assessed with gel electrophoresis.

Library Preparation was performed by Edinburgh Genetics using egSEQ Enzymatic DNA Library Preparation (Edinburgh Genetics, n.d.). This involved DNA fragmentation and selection of insert sizes between 300-400bp. End repair and adenine base addition (A-tailing) enabled adapter ligation before DNA purification and amplification via PCR. Circularisation was performed to ensure the library was suitable for DNBSEQ technology. Whole genome sequencing was carried out using MGI DNBseq-T7 on paired-end reads of length 150bp. Around 22Gb of sequence data was supplied per sample, giving 50X genome coverage. Data quality control and filtering was performed using soapnuke software (Chen et al. 2018).

### Trimming and Mapping

Sequencing quality was evaluated using fastQC (v0.11.9) (Andrews 2010) and multiQC (v1.14) (Ewels et al. 2016). Filtering was carried out using NGSQCToolkit IlluQC.pl (Patel and Jain 2012) where reads were removed if more than 70% of bases had a Phred below Q20. Reads were trimmed using trimmomatic (v0.36) (Bolger et al. 2014) according to the following thresholds: LEADING:10, TRAILING:10, SLIDINGWINDOW:4:20, and MINLEN:70.

Sequences for the 324 genotyped individuals were mapped to the Darwin Tree of Life (DToL) reference genome of *Alnus glutinosa* (Assembly version GCA_958979055.1, available from https://www.ebi.ac.uk/ena/browser/view/GCA_958979055 (Darwin Tree of Life Project, n.d.) using bwa mem (v0.7.17) (Li and Durbin 2009). Samtools (v1.10) (Li et al. 2009) -fixmate and -markup were used to mark and remove PCR duplicates. Samtools -view generated BAM format files filtering for a mapping quality above 20. Alignment quality was assessed using SamTools and QualiMap (v2.2.1) (García-Alcalde et al. 2012). Additional Qualimap metrics (mapping quality and genome-wide coverage distribution per sample) were assessed.

### SNP calling and filtering

Variant calling was performed using GATK HaplotypeCaller (v4.2.6.1) (Poplin et al. 2017). GATK GenomicsDBImport imported the 324 samples into a single database (batch size of 20). Joint variant calling was performed using GATK GenotypeGVCFs (DePristo et al. 2011; Van Der Auwera. GA 2020). Variants were counted using GATK CountVariants prior to filtering for indels and mixed type variants using GATK SelectVariants (-select-type-to-include=SNP).

SNP allelic balance was plotted for a 30% subset of SNPs and chromosome-by-chromosome for each individual in Rstudio (v4.2.2) (R Core Team 2026). Allele balance plots identified two triploids (Figures S1c and S1d) and one sample with ambiguous ploidy (Figure S1g). The sample import step was repeated on 321 samples following removal of these individuals. Two samples showing ambiguous allele balance (Figures S1e & S1f), and a trisomic individual (Figure S2) were retained pending further investigation.

SNPs underwent three rounds of filtering: GATK, vcftools (v0.1.16) (Danecek et al. 2011), and plink2 (v2.0-20220603) (Chang et al. 2015). GATK was used to flag and remove SNPs failing the following thresholds: quality by depth (QD) < 2.0, quality (QUAL) <30, StrandOddsRatio (SOR) >2.5, FisherStrand (FS) >60.0, root-mean-square Mapping Quality (MQ) <50.0, Mapping Quality Rank Sum (MQRankSum) <-2.5 and >2.5, Read Position Rank Sum (ReadPosRankSum) >2.0 and <-2.0. Bi-allelic SNPs were selected using bcftools view with the -m2 and -M2 options (v1.16) (Li, 2011). Vcftools was used to filter SNPs with missing rates >15%, minor allele count (mac) <3, minor allele frequency (maf) <0.04, minimum quality (minQ) <30, minimum depth (minDP) <15, and maximum depth (maxDP) >100.

Variants strongly deviating from Hardy-Weinberg equilibrium (i.e. p-values <1^-50^), and individuals with missing rates >10% were removed using plink2, which saw the removal of two samples. This left 7,375,952 high quality SNPs. Density of these SNPs along the genome was assessed using the *CMplot* R package (Yin et al. 2021) (Figure S3).

### Ploidy Analysis and Outlier Identification

To investigate ambiguous allelic balances, mapping quality and coverage were assessed for all 324 individuals following the first variant call (see *SNP calling and filtering*). SNP quality filtering on all 324 individuals was repeated, but without the individual-level missing data filtering step in plink2 to retain all individuals (see *SNP calling and filtering).* Principal Components Analysis (PCA) was performed in plink2. Missingness (F_MISS) and inbreeding coefficient (F) was extracted using vcftools with the --missing-indv and --het flags. Sequences were evaluated for DNA contamination using blast (v2.11.0) (Camacho et al. 2009) and blobplots (v4.3.3) (Laetsch and Blaxter 2017) with no contaminants identified.

Assessments of the metrics above identified two outliers matching those with ambiguous allele balance (Figures 1f, S2f & S2g, see *SNP calling and filtering*). For the 321 individuals included in the second variant call (which reduced to 319 following filtering, see *SNP calling and filtering)*, these two outliers were removed. SNPs in linkage were pruned using plink2 – indep-pairwise with an r^2^ threshold of 0.4 within a 50bp window, shifting 4bp along at a time, leaving ∼1.49M SNPs in approximate linkage equilibrium. Technical replicates were identified using a kinship matrix calculated using plink2 (-–make-king) and plotted as a heatmap using ggplot2 (v4.0.2) (Villanueva and Chen 2019). These were removed prior to association testing.

**Figure 1.**
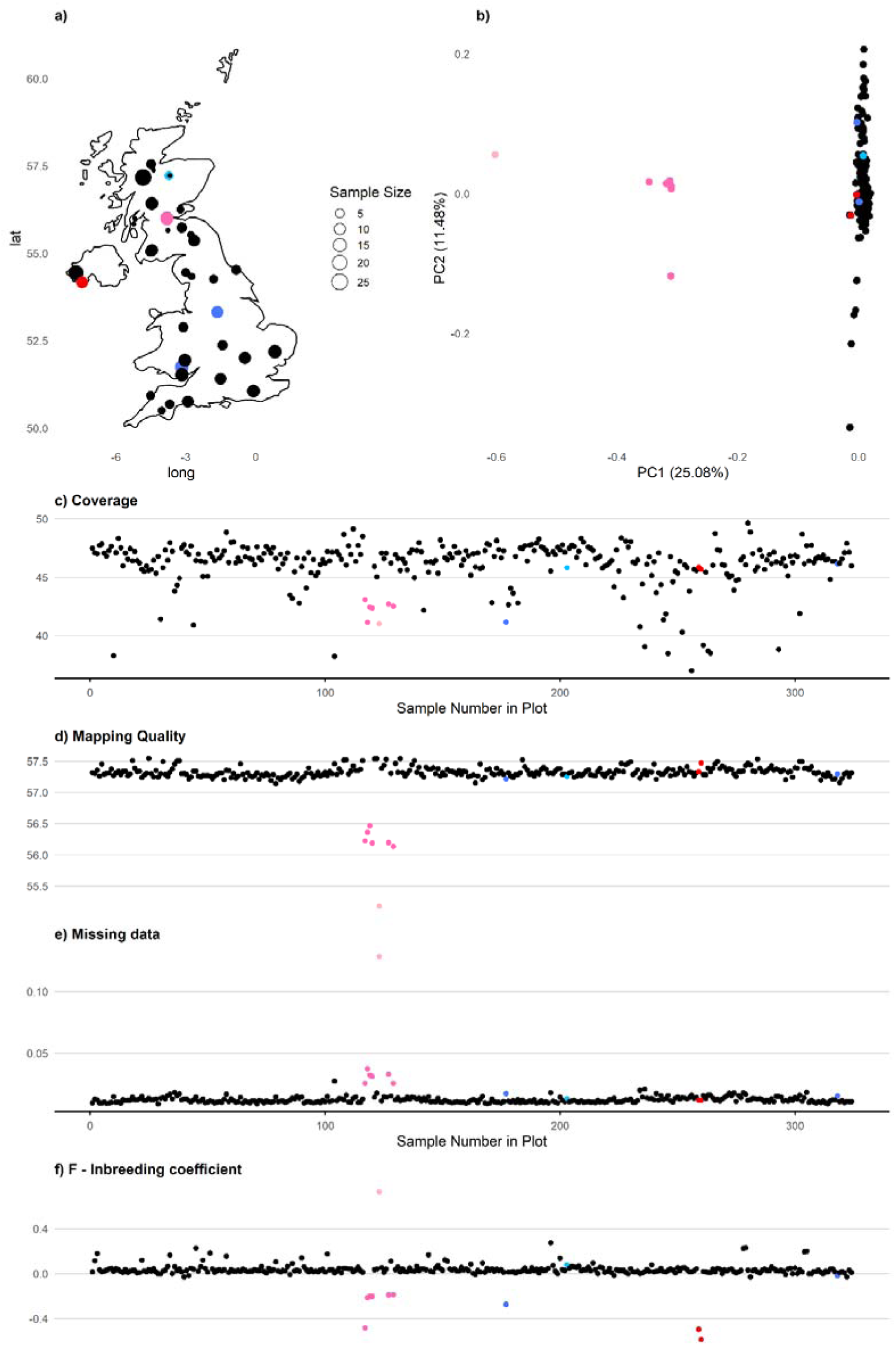
Location and characteristics of 324 UK *Alnus glutinosa* individuals in 44 populations. **a** shows the location and sample size of each population. **b** shows a PCA. **c-f** show quality characteristics per sample including **c,** coverage **d,** mapping quality, **e** missing data, and **f,** inbreeding coefficient. Dark blue indicates triploids, light plink for 0550 (ambiguous ploidy likely caused by high missingness), red for samples for which ploidy could not be determined, dark pink for diploid hybrids, and light blue for the trisomic sample.

Ancestry proportions and population genetic structure was inferred using fastSTRUCTURE (v1.0) (Raj et al. 2014) and principal component analysis (PCA). fastSTRUCTURE was run with 1.49M LD-filtered SNPs from K1 to K25 with 5 cross validation tests. fastSTRUCTURE’s ‘chooseK’ and ‘distruct’ scripts were used to estimate the best K to explain the components of the data and to plot admixture proportions by individual, respectively. Kinship and fastSTRUCTURE identified six samples within population 899279 as different from the rest of the dataset, matching outliers on PCA plots, and for mapping quality, missing rates, and inbreeding coefficient. The six outlier individuals were removed prior to association testing.

### Population Structure

For the remaining 304 individuals, PCA, kinship and fastSTRUCTURE analyses were repeated (as described in *Ploidy Analysis and Outlier Identification* section above). Estimates of population genetic diversity were calculated using Stacks (v2.41) (Catchen et al. 2013) on a 10% (148,931) subset of LD-filtered SNPs. Estimates were regressed onto latitude using simple linear regression models in Rstudio using the lm() function from the stats package. Mean pairwise F_st_ was calculated for each population and regressed against latitude using a simple linear regression model. Statistical significance was assessed using F-tests.

### Collating Environmental Data

Nineteen 1 km-resolution temperature and precipitation climatologies were downloaded from the Climatologies at High Resolution for the Earth’s Land Surface Areas (CHELSA) WordClim global database (Karger et al. 2017) version 2.1 and imported into Rstudio using the *Raster* package (v3.6.32) (Hijmans 2025). Variables are annual averages between 1981-2010 and reflected annual trends, seasonal variation, and climate extremes (Table S3).

Four additional 1 km-resolution datasets for 1991-2010 were downloaded from the MetOffice HadUK-Grid dataset (Met Office et al. 2024) and added to the raster stack with consistent coordinate reference system assignment. These were annual mean frost days, average snow days, average wind speed and sunshine duration (Table S3). Coordinate locations of alder populations were used to extract values for each environmental variable using raster::extract().

Altitude data were downloaded from https://biogeo.ucdavis.edu/data/diva/alt/GBR_alt.zip, and raster was used to extract slope, aspect, roughness, topographic position index (TPI), and terrain ruggedness index (TRI). Aspect was transformed from radians into a continuous variable using cos() and sin(), dividing it into northing and easting respectively. This resulted in 29 environmental variables. The CHELSA variables were offset prior to analysis.

Variable correlations were visualised using removeCollinarity(), within the *virtualSpecies* R package, with a Pearson’s coefficient of 0.7 (Figure S8a) (Leroy et al. 2016). A heatmap of pairwise correlation coefficients was created using the *corrr* (Kuhn et al. 2016), *ggcorrplot (Kassambara 2022)*, and *corrplot* packages (Wei and Simko 2010) (Figure S8b). To remove correlated variables, an R script iteratively removed one variable from each highly correlated pair (r >=0.7) (Carleial et al. 2025). Variables in high-correlation pairs were evaluated based on their mean absolute correlation to all other variables, and that with the highest mean correlation was removed. This was repeated until no pairwise correlations exceeded 0.7, resulting in 11 variables (Table S3). These were standardised to have mean of 0 and standard deviation of 1 across all 304 individuals.

### Genome-Environment Association

Associations between SNPs and environmental gradients were investigated using latent factor mixed models (LFMMs) implemented in the Landscape and Ecological Association (*LEA*) R package (Caye et al. 2019; Frichot et al. 2026). The frequentist least-squares lfmm2() was chosen, as opposed to the Bayesian lfmm(), because the former has been shown to produce less false positives (Caye et al. 2019). Missing genotypes were imputed using BEAGLE (v5.4) (Browning et al. 2018). lfmm2() requires a user set K value of latent factors (number of ancestral populations/sources of hidden population structure). To determine an appropriate K value, sparse non-negative matrix factorisation (SNMF) was run within the LEA package. This uses cross-validation to test the appropriateness of each K value at explaining variation in genotype data (Frichot et al. 2014, 2026). SNMF was run with K values from 1 to 10 and entropy = TRUE for 10 repetitions on a dataset of 1.49 million unlinked SNPs. Cross-entropy was plotted for each K, with the lowest value indicating the K which best explains structure in the genotype data (Frichot et al. 2014; Alexander and Lange 2011). PCA was used to complement the SNMF analysis, and was run within the *LEA* package. Eigenvalues were plotted to identify the number of components explaining meaningful population structure. SNMF suggested K2 or K4 best explained the genotypic variation (lowest cross entropy, Figure 2d), and PCA indicated two genetic/ancestral populations according to Cattel’s rule (elbow of the plot) (Frichot et al. 2026).

**Figure 2.**
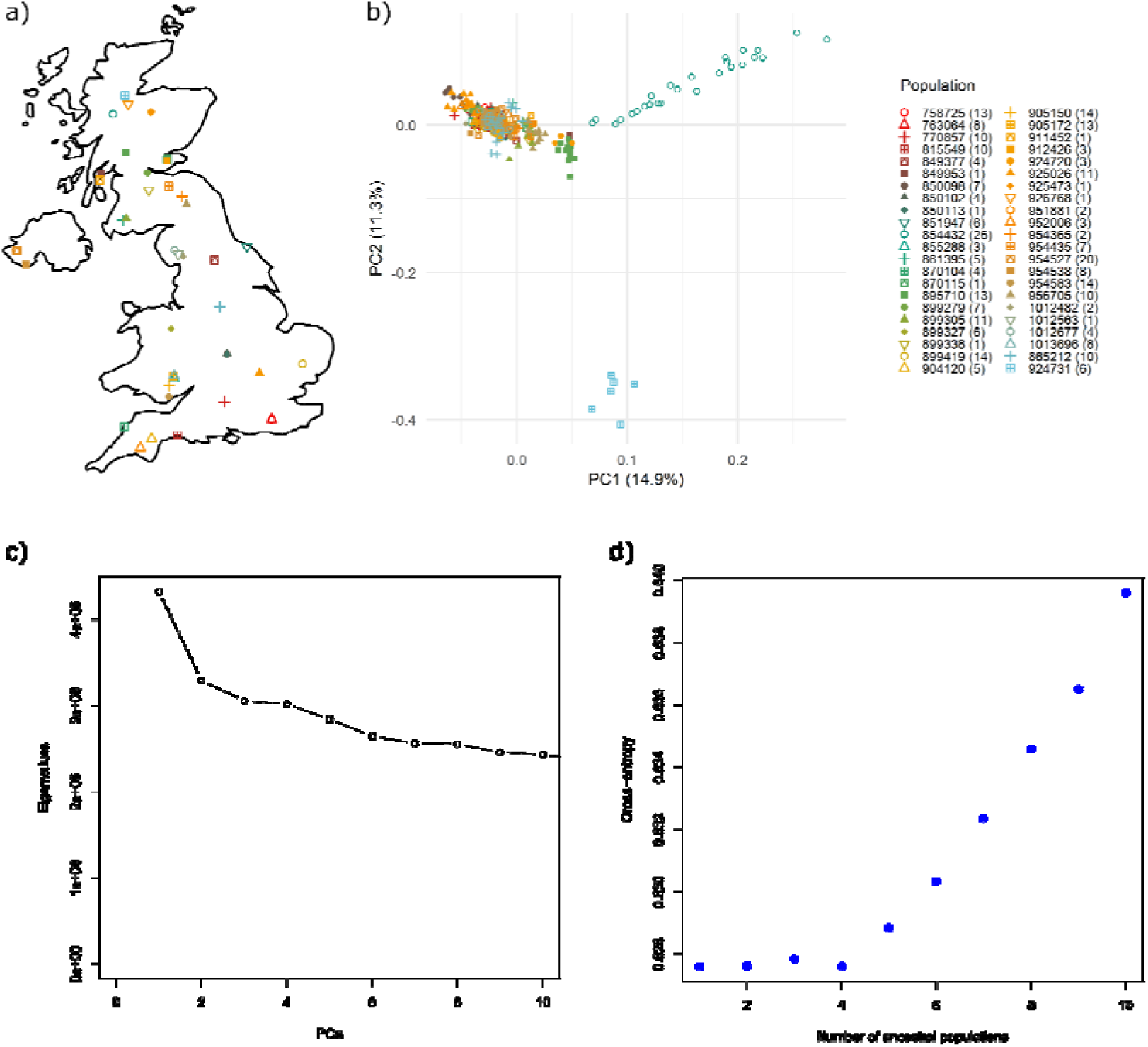
Population Structure of 304 diploid *Alnus glutinosa* individuals. **a** shows the location of each population. **b** shows a PCA. The colour scheme is consistent across panels a and b. **c** shows a scree-plot of eigenvalues following PCA, where the elbow of the plot represents the optimum K value. **d** shows SNMF analysis, where lower cross-entropy indicates better K value. Analysis shown in c and d was performed using the *LEA* R package.

After identification of the appropriate K, lfmm2() was fitted with ∼7.4M genome-wide SNPs from 304 diploid individuals and 11 uncorrelated environmental variables, with the effect.size and linear arguments set to TRUE. P-values for the lfmm2() associations were generated using the lfmm2.test() function in LEA with linear = TRUE and genomic.control = TRUE (Frichot et al. 2026). Models were re-run at K1, K3 and K4.

### Manhattan plots and annotation

P-values were plotted as Manhattan plots using the *topR* package (Juliusdottir 2023). To control for false positives, a false discovery rate (FDR) was performed by transforming p-values into q-values using the R package *qvalue* (Bass et al., 2023) and applying a q-value threshold of 0.05. For comparison, two more stringent thresholds were included in the Manhattan plots: 1) Bonferroni correction: dividing 0.05 by the number of SNPs; and 2) modified Bonferroni: dividing 0.05 by the number of haplotype blocks in the genome, determined using plink2 --blocks function. Genomic inflation was assessed using Q-Qplots made in *GWASTools* v1.44.0 (Gogarten et al. 2012). Population-level allele frequencies of significantly associated SNPs according to the FDR threshold were extracted using plink2 and plotted against local conditions of each population for the respective variable of association.

Genes near the SNPs significantly associated with the environment were identified using a custom R function (Carleial et al. 2025) and the *A. glutinosa* gene annotation file available at https://rapid.ensembl.org/Alnus_glutinosa_GCA_958979055.1/Info/Index?db=core;g=ENSGUIG00005018309;r=6:20492624-20498837;t=ENSGUIT00005024060 (Darwin Tree of Life Project).

### Variable contribution to genomic signal

The genomic signal of each environmental variable was estimated using the SNP effect size matrix (β) generated in the LFMM2 above, as described in (Carleial et al. 2025). Briefly, a covariance matrix of genetic effects between environments (Σ_β) was generated. For each variable, the total genomic variance (var_total) was determined as the diagonal of Σβ, representing the total variance in SNP effect sizes. Redundancy - or the variation in β explained by more than one variable - was determined by the difference between total and conditional variance (var_cond), where conditional variance is the inverse of the diagonal element of the matrix. Relative uniqueness was calculated as the fraction of independent genomic signal not shared with other variables (var_cond / var_total). These metrics allow partitioning of associations into shared and independent components, facilitating identification of dominant environmental drivers of genomic variation.

### Predicting genetic change

Future, projected annual averages were extracted for each CHELSA climatology for a period of 2071-2100 based on the worst-case climate change scenario (Representative Concentration Pathway (RCP) 8.5) from the UKESM1-0-LL model. Annual mean wind speed between 2071-2081 for RCP8.5 at 5-km resolution was extracted from theMetOffice UK Climate Projections (UKPC) database (Met Office) available at https://ukclimateprojections-ui.metoffice.gov.uk/ui/home. The number of days with snow lying was not available on the UKPC database so could not be included. Annual mean frost days was also not available through UKPC databases, but a similar dataset (annual number of days where the minimum daily temperature is below 0°C) was available via the MetOffice climate data portal (Met Office 2023), available at https://climatedataportal.metoffice.gov.uk/datasets/TheMetOffice::annual-count-of-frost-days-projections-12km/about). Projected frost data at 12-km resolution was extracted for a 21-year time period centered around when the annual mean temperature reaches 4.0°C under RCP8.5. Terrain data (see *Environmental Variable Selection* section) was added, and the matrix was subset to predictors retained in the original lfmm2() following removal of highly correlated variables, resulting in 10 final variables.

To ensure compatibility between current (1980/1991 -2010, see *Collating Environmental Data* section) and future climate analyses, the same ten environmental variables were used for both time points. Because annual mean snow days was unavailable for future climate projections, it was removed from the current environment matrix, which was then re-standardised, and the lfmm2() was re-run. The future dataset was standardised using the scaling parameters estimated from the current climate dataset.

### Genomic Offset - LEA

The genetic.offset() function in the LEA package was implemented to predict the degree of genetic change required for populations to remain adapted to future conditions. Following Gain et al., (2023), this function uses the covariance matrix of SNP effect sizes estimated from lfmm2() to compute the quadratic distance (geometric genomic offset) between current and future conditions for each location. This produces a genetically-weighted measure of maladaptation, where the contribution of each environmental variable is weighted by its effect on allele frequencies across loci. This was run on all 1.49 million unlinked SNPs.

We tested the assumption that larger genomic offsets will be observed for populations with greater disparity between current and future climates by quantifying differences between time points using PCA. We used the first five components, which explained over 78% of variation. Environmental dissimilarity was calculated as the Euclidean distance between current and future coordinates for each population within the multidimensional PC space.

Linear models were performed within the *stats* base R package to assess trends between genetic offset, environmental dissimilarity, latitude and elevation i.e. GeneticOffset∼ EucDistance. A multiple regression was performed to investigate the contribution of each metric on genetic offset after accounting for the other i.e. GeneticOffset ∼ EucDistance + Latitude. P-values were estimated with F-tests.

### AlleleShift

Genomic offset estimates were compared to estimates of maladaptation using the *AlleleShift* (Kindt 2021b) R package. This method attempts to predict the change in allele frequency of environment-associated SNPs required for adaptation to a different set of environmental conditions (present or future). The AlleleShift pipeline is comprehensively described in Kindt (Kindt 2021a). Briefly, the *vcfR* (Knaus and Grünwald 2017) and *adegenet* (Jombart 2008) packages were used to generate allele counts per population for 1,364 significant SNPs, and the top 10,000 SNPs with the lowest p-values. RDA models were trained on each SNP set to learn the relationship between allele frequencies and present-day environmental variables (See *Extracting future variables*). Models were calibrated with a GAM to eliminate negative allele frequencies. The calibrated models were used to predict the optimum allele frequencies expected under current, and future environmental conditions for each population. The allele shift (proxy of mal-adaptedness) was calculated as the absolute difference between the predicted, optimum frequency and the current, baseline frequency. Regressions were repeated as above (see *Genomic Offset - LEA*) for mean shift, latitude, and allele shift environmental dissimilarity.

## Results

### Data generation

Average coverage per base following alignment of each of the 324 samples to the reference genome was 45.0X and 46.1X for Samtools and Qualimap respectively. The mean mapping quality score across samples was 57.3. A total of 54,281,745 raw variants were identified across all 324 individuals, of which 43,088,288 were single nucleotide polymorphisms (SNPs).

### Screening for polyploidy, polysomy and outliers

Allele balance showed a distribution consistent with diploidy in 318 out of 324 individuals (Figure S1a and S1b). Two individuals showed a triploid pattern and were not retained in further analyses (Figure S1c and S1d). An individual with trisomy was identified (Figure S2), the triplicated chromosome was removed, and retained in subsequent analysis.

Three samples exhibited ambiguous allele balance for which ploidy was hard to determine (Figure S1e, S1f and S1g). For two of these samples from population 954538 (0851 and 0852), the location of peaks in histograms of the number of reads supporting each allele suggested either a high ploidy level, cross contamination, or technical artefacts. We removed 0852, but retained two ambiguous samples pending further investigation.

Following the removal of polyploids, variant calling on 321 individuals resulted in ∼54 million variants of which ∼43 million were SNPs. After hard filtering, ∼24 million SNPs remained which were reduced to ∼21 million after filtering for bi-allelic variants. VCFtools filtering removed 14 million SNPs and plink filtered out a further 88K (see Methods section *SNP calling and filtering* for thresholds). Two individuals were removed due to high SNP missing rates (0550 and 0684), one of which was an individual with ambiguous ploidy identified above, and the other the individual with putative trisomy. Following removal of technical replicates and the remaining individual for which ploidy was ambiguous (0851, Figure S1f), 311 samples from 44 populations remained.

### Population structure analysis

PCA of the SNP data following the first variant call (see *SNP calling and filtering*), differentiated six individuals across PC1, which explains 25.01% of variance (Figure 1b). Further investigation found their SNP calls had low average mapping quality, low inbreeding coefficients, and high proportion of missing data (Figure 1d, 1e, 1f). These six individuals were diploid, represented six out of thirteen individuals in population 899279, and were closely related to each other (Figures S4b & S5a). This pattern suggests natural hybrids, or accidental mixing of samples in the laboratory. Population admixture estimated with fastSTRUCTURE including all individuals identified K2 as the model which maximises the marginal likelihood, and K5 best at explaining structure in the data. After re-running fastSTRUCTURE without the six outliers, these values were reduced to K1 and K4, respectively.

Following removal of the six outliers, PCA separated two northern populations (Figure 2b). Samples from Scottish population 854432 splintered across PC1, which explained 14.9% of variation, and Scottish population 924731 was separated by PC2, which explained 11.3% of variation (Figure 2b). These populations were separate clusters in admixture analyses above K=2 (Figure S5). The fact that population 854432 has the largest sample size (n = 26, compared with a mean of 6.44 ± 4.77 SD across all other populations) may have contributed to this pattern, by capturing more within-population diversity and increasing its contribution to the PCA covariance matrix. The population has relatively low average pairwise F_ST_ (Figure S6h), which taken alone indicates little differentiation from other populations. Population 924731 (n = 6) has a slightly negative inbreeding coefficient (F_IS_) of -0.05 compared to a mean of 0.006 across all other populations (Figure S6g, note the small y-axis scale showing F_IS_ values were close to zero in all populations); this marginal heterozygote excess may reflect outbreeding, contributing to the separation observed during PCA.

Population latitude showed a significant, positive relationship with mean frequency of the most frequent allele (P) and a significant, negative association with expected heterozygosity (Exp_Het, Figure 7S; Table S2), implying reduced genetic diversity in northern populations. Mean pairwise F_st_ increased significantly with latitude (F□,□□ = 7.70, p = 0.008, R^2^ = 0.15) suggesting northern populations were generally more genetically differentiated from other populations (Figure S6h).

Aside from 854432 and 924721, remaining samples form one cluster in the PCA indicating little overall population structure (Figure 2b). This was corroborated by *LEA* SNMF analysis indicating low values of K (K1, K2 or K4) as optimal to explain variation in the genotypic data (Figure 2d). *LEA* PCA complemented this, with 2 ancestral components best at explaining structure according to Cattel’s rule (Frichot et al. 2026, Figure 2c). FastStructure *ChooseK.py* reported K1 as the best model to explain the data, and four detectable ancestral components indicating little genetic structure. We used K=2 to control for population structure in the GEA (see below).

### Genome-Environment Association

7.38 million SNPs from 304 individuals in 44 populations were assessed for association to 11 environmental variables at K2. In total, 1,364 SNPs were identified as significantly associated with the environment according to the FDR threshold. The majority of these (1,269) were associated with the average number of snow days (Figures 3 and S12), for which genomic inflation was high (Figure S11). Significant SNPs were identified in discrete peaks across many chromosomes (Figure 3). An additional 36 SNPs were associated with frost, 26 with TPI, 13 with isothermality, 13 with mean temperature of the wettest quarter, five with slope, and two with precipitation of the driest month (Figures 3 and S12). Two SNPs (11:27300481 and 11:27300569) were significantly associated with precipitation of the driest month, mean temperature of the wettest quarter, and isothermality. Four SNPs (1:18857259, 1:18857282, 1:18857301, 1:18857307) were found significantly associated with both average number of snow days and number of frost days. No SNPs were found associated with northing, mean temperature of the driest quarter, easting or wind speed (Figures S10 and S12). The same variables showed significantly associated SNPs irrespective of the K value used (Figure S12).

**Figure 3.**
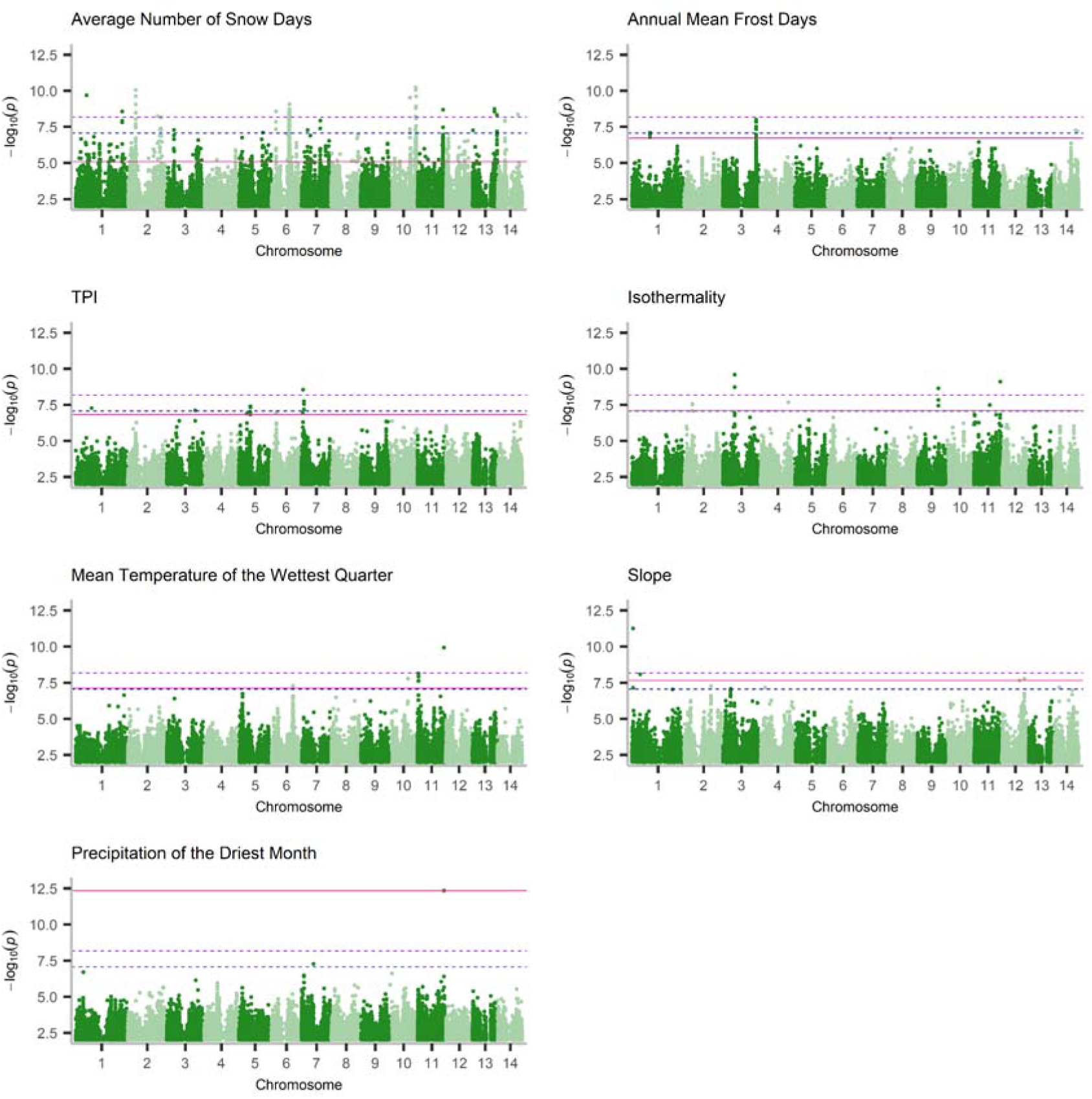
Genome-environment association study in UK *Alnus glutinosa.* Each panel shows a Manhattan plot for an environmental variable, with -log10 p-values on the y-axis and position on the x-axis. Horizontal lines indicate FDR (pink), modified-Bonferroni (blue), and Bonferroni (purple) significance thresholds. Variables shown are those with significance according to the FDR threshold. Latent Factor Mixed Models were run on 7.38 million SNPs at K2 on 304 individuals and 11 environmental variables. Plots for variables without SNPs above the FDR significance threshold are shown in Figure S10.

Population-level allele frequencies of the most significantly associated loci show clines across environmental gradients for number of snow days and frost days (Figure S13). Trends for other variables are skewed by outlier populations, most noticeably for slope where the allele is fixed in population 926768, but absent in nearly all other populations (Figure S13). This calls into question the reliability of ‘significant’ SNPs, as the strength of the association appears to be driven by extreme values in the variable’s distribution (Figure S9). This prompted further analyses exploring the relationship between environmental variables and variation in genome-wide SNP effect sizes from lfmm2(). This approach is best suited for polygenic traits and avoids overreliance on significant loci (Carleial et al. 2025).

To capture small-effect SNPs typical of polygenic local adaptation, the contribution of each environmental variable to genomic variation across all 7.38 million genome-wide SNPs was assessed. The greatest total variance in SNP effect sizes was observed for mean temperature of the wettest quarter, contributing 15.6% of total variance, followed by annual mean frost days and precipitation of the driest month responsible for 13.9% and 12.7% of total variance respectively (Figure 4a). These variables also exhibit the greatest redundancy (Figure 4a), suggesting that collinearity persists among environmental predictors despite the prior removal of highly correlated variables (|r| > 0.7) before GEA.

**Figure 4.**
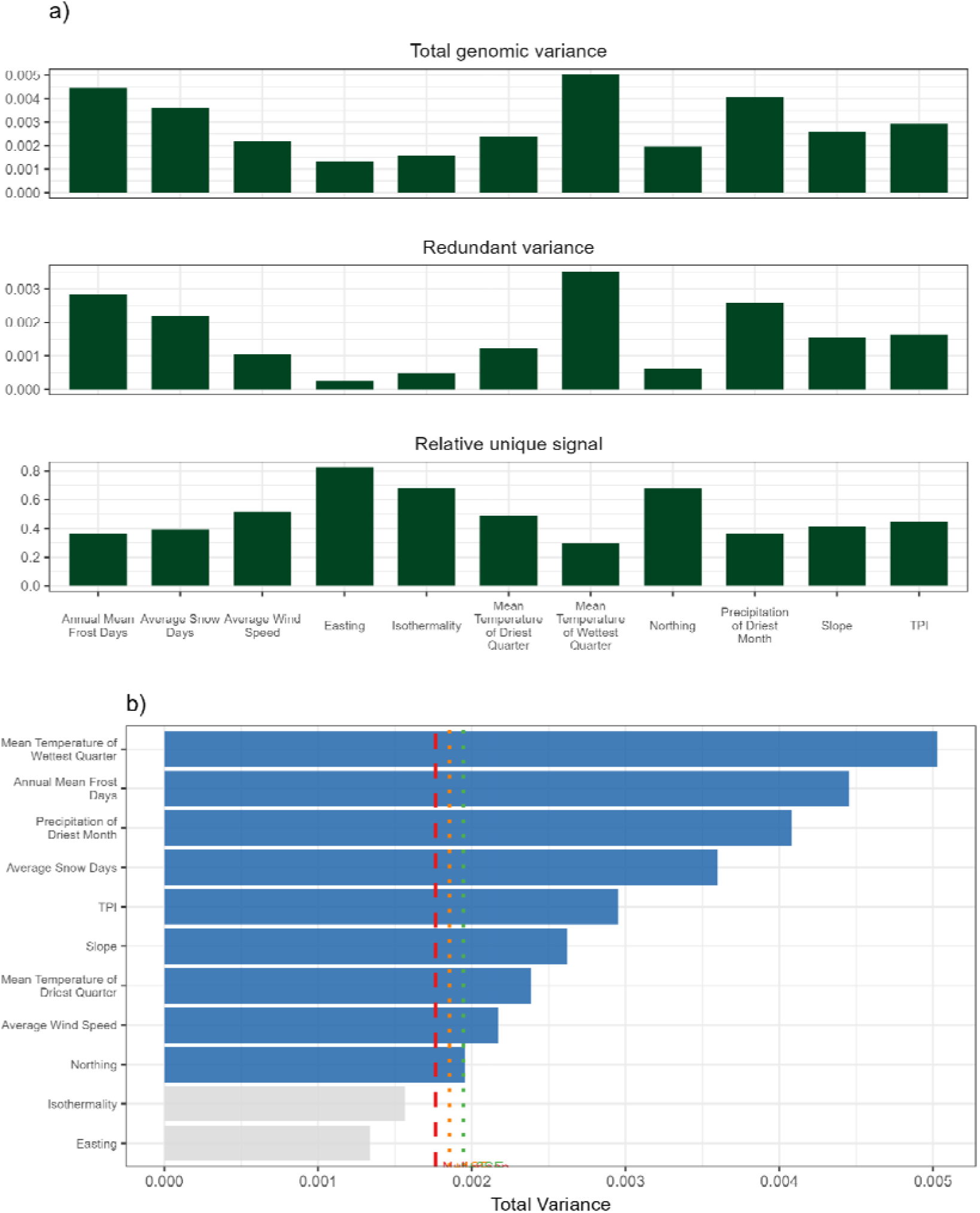
Contribution of environmental variables to genome-wide genetic variation. **a,** the total, shared and unique variation in SNP effect sizes attributed to each variable following covariance assessment of the SNP effect size matrix (see Methods, *Variable contribution to genomic signal)*. **b,** the total variance of each variable compared to the mean (red dashed line), +1SE (orange dashed line), and +2SE (green dashed line) obtained from ten randomly generated dummy variables to assess deviation from a null expectation. Variables with a total variance >2SE from the mean were deemed genuine (blue bars), whilst those lower than the null were indistinguishable from noise (grey bars). Latent Factor Mixed Models were run on 7.38 million SNPs at K2 on 304 individuals and 11 environmental variables.

After accounting for conditional variance, easting, isothermality and northing contributed the greatest proportion of independent genomic variation, accounting for 15%, 12.5% and 12.4% of the total independent genomic signal (Figure 4a). These variables also had the lowest total genomic signal suggesting they contribute in a minor but unique way to SNP effect size. Ten randomly generated dummy variables contributed a null average 4.98% ± 0.735 SD of total variance in SNP effect size. The majority of nine variables exceeded this null by more than two standard errors suggesting genuine genomic signals (Figure 4b). Easting and isothermality fell below the null suggesting their signal was indistinguishable from noise. This warrants caution in interpreting SNPs significantly associated with these variables.

### Predicting genetic change required for adaptation

Northern populations exhibit a significantly greater difference between present-day, observed allele frequencies and predicted, optimum frequencies in current climates (Figure S14b). This was driven by 924720 and 925473 when ∼1.3K significant SNPs and the top 10K LFMM SNPs ordered by p-value were used for estimations (Figure S14). These populations are located geographically close together (Figure 2a) and are outliers for average number of snow days, which had considerably more significant SNPs (Figure S9 and S12). This suggests loci associated with cold environments contribute disproportionally to the observed pattern of present-day maladaptation. Given high inflation observed for this variable, which lowers the confidence in the identified significant SNPs, we trust *AlleleShift*-estimated maladaptation using SNP subsets less than the *LEA* genomic offset approach using genome-wide markers.

Climate dissimilarity between current and predicted environmental conditions in 2071-2100 under the now defunct RCP8.5 (calculated as the Euclidean distance in PCA space between timepoints for each population, Figure S15) showed a significant relationship with latitude (F□,□□ = 6.76, p = 0.013, R² = 0.14, Figure 5a & 5e). Northern populations are predicted to experience significantly greater climate shifts than southern locations (Figure 5a & 5e). Exceptions included population 924720 and 925473 in Scotland for which predicted climate shifts were lower than may be expected given their latitudes (Figure 5a & 5e). These were the same populations identified with the greatest mal-adaptation to current local climates (see above, Figure S14).

**Figure 5.**
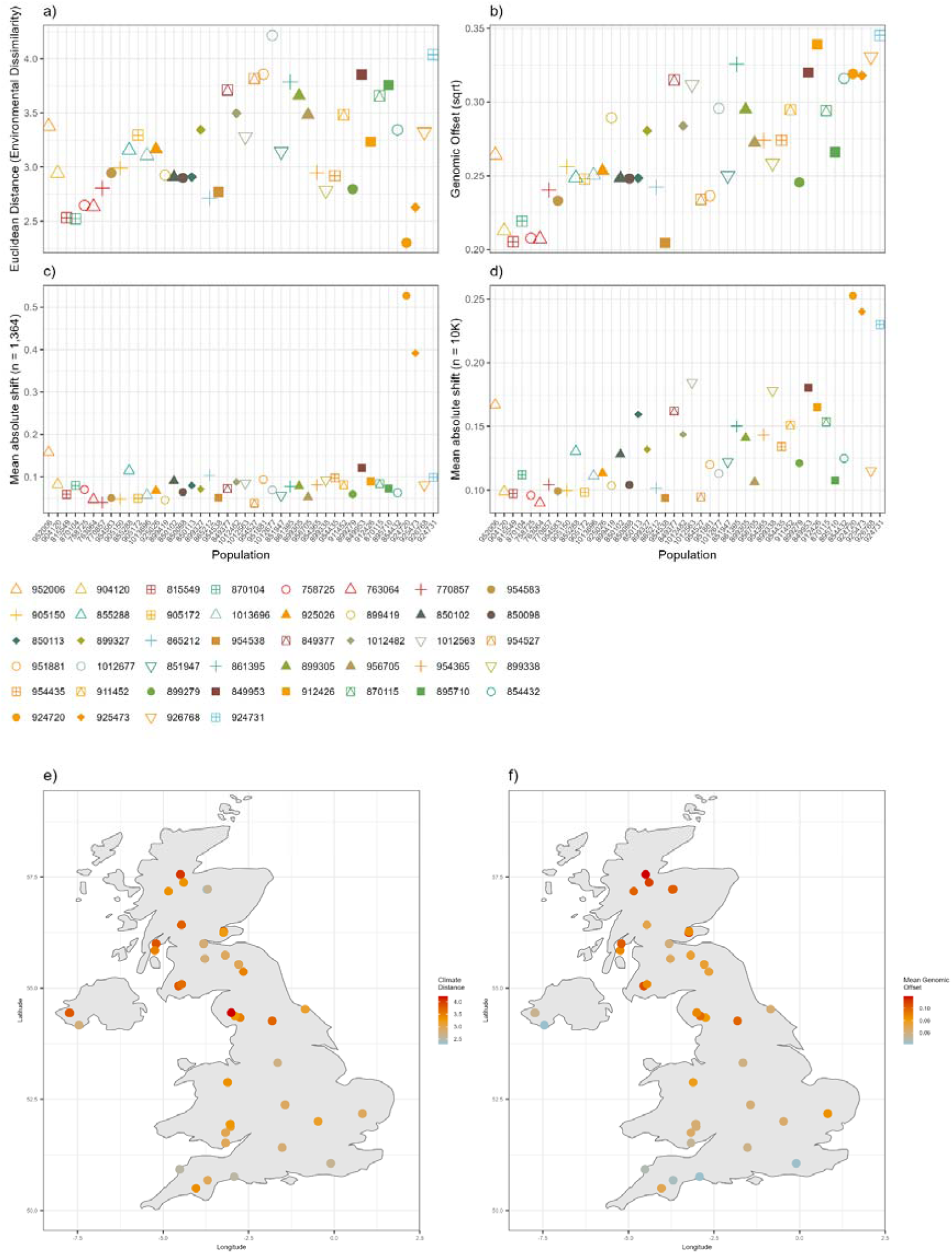
Predicted environmental and genetic change across 44 UK *Alnus glutinosa* populations. **a,** Environmental dissimilarity between current and future local climates calculated as Euclidean distance in PCA space. **b,** Genomic offset required for populations to remain adapted to future climates, estimated using LEA on 1.49 million unlinked SNPs. **c,** Mean allele frequency shift required for climate-associated SNPs to reach future optimum population frequencies, estimated using AlleleShift on 1,364 FDR-significant SNPs. **d,** Mean allele frequency shift estimated using the 10,000 lowest *p*-value SNPs from LFMM2. In **a–d**, populations are ordered by latitude. **e,** Population locations coloured by environmental dissimilarity and **f,** by genomic offset. Future climate predictions represent the RCP8.5 scenario for 2071–2100.

The *LEA* R package was used to estimate the genetic change required for each population to be adapted to future local climates using 1.49 million LD-filtered SNPs (henceforth called ‘genomic offset’). A significant positive relationship was identified between genomic offset and climate dissimilarity (F□,□□= 12.91, p < 0.001, R² = 0.24, Figure S16a), and between genomic offset and latitude (F□,□□ = 49.43, p < 0.001, R² = 0.54, Figure 5b and 5f). This suggests that northern populations need to undergo larger genetic change than southern ones to match the optimum allele frequencies predicted for their future local climates. No significant trends were identified between elevation and either climate dissimilarity or genomic offset (Figure S17).

When both latitude and climate dissimilarity were included in a multiple regression, they jointly explained 59% of the variance in genomic offset. Latitude was a better predictor of variation in genomic offset (F□,□□ = 35.94, p < 0.001) than climate dissimilarity (F□,□□ = 5.21, p = 0.028). The inclusion of climate dissimilarity to the model only increased R^2^ by 5% over the model with latitude alone.

A similar pattern was observed for *AlleleShift*-estimated genetic maladaptation using 10K climate-associated SNPs (henceforth called ‘allele shift (10K SNPs)’); the shift in allele frequency required for present day populations to match predicted optimum allele frequencies in future climates increased with latitude (F□,□□ = 20.55, p < 0.001, Figure 5d). Contrasting *LEA* genomic offset estimates, the allele shift (10K SNPs) estimates were not significantly associated with climate dissimilarity (F□,□□ = 0.33, p = 0.57, Figure S16b).

When only ∼1.3K FDR-significant SNPs were used in *AlleleShift*, the relationship between maladaptation and latitude disappears (Figure 5c). Two outlying populations become apparent; 924720 and 925473, mirroring patterns of predicted genetic change for current conditions using the same SNP set (Figure S14). The genetic change predicted for adaptation to both current and future climates is higher for these locations using the *AlleleShift* approach on a reduced SNP set, but these same sites are predicted to experience the least environmental change (Figure 5a, 5c, and S14). This highlights the lack of reliability in estimates of genomic offset using reduced, significant SNP sets which don’t account for effect-sizes or correlations among environmental variables.

*Alleleshift* and *LEA* genomic offset estimates were somewhat correlated. Allele shifts estimated using only significant GEA SNPs show a Spearman’s rank correlation coefficient of 0.46 with *LEA* estimates (Figure S18a). The correlation increases to 0.59 when the top 10k SNPs are used instead. Estimates from the same model and SNP sets showed high correlations suggesting the model and SNPs utilised in the analysis have a large impact on the estimates of maladaptation. Predictions based on larger SNP sets showed stronger correlations between predicted genetic change and climate dissimilarity than predictions based on smaller SNP sets (0.5 for genomic offset, 0.27 for AS future 10k, 0.07 for AS future 1k, Figure S18a). The genome-wide *LEA* estimates also showed a greater correlation with latitude (0.72) than the *Alleleshift* estimates (0.58 for AS future 10k, 0.38 for AS future 1k). Removing the two outlier populations had little impact on the genome-wide estimated correlations, but strengthened the relationship between allele shifts and climate distance (Figure S18).

## Discussion

We find that UK native *Alnus glutinosa* populations have low genetic differentiation consistent with high gene flow through wind- and water-dispersed pollen and seed. Genetic diversity declined and differentiation increased with latitude, consistent with genetic drift at the northern edge of post-glacial recolonisation. Triploids and the first reported case of trisomy in this species were also detected. Genome-wide approaches to GEAs identified temperature- and precipitation-related variables as associated with the greatest genetic variation. Analysis of only statistically significant GEA SNPs gave contrasting estimates of genomic offset compared to genome-wide SNPs, and appeared less reliable owing to outlying population-level allele frequencies in populations with extreme local environmental values. Genomic offset identified northern populations as the most genetically vulnerable to both current and future climates highlighting the potential role of assisted gene flow from south-to-north. Seed sourcing for restoration programmes should prioritise populations from environments with temperature and rainfall similar to those at the planting site, thereby promoting adaptive capacity and survival by matching allele frequencies with local climate.

### Hybrids and Polyploids

We found six samples separating during PCA with a high proportion of missing data, and low mapping quality and inbreeding coefficient, indicating greater heterozygosity consistent with natural hybridisation or laboratory contamination. Four other *Alnus* species are present in the UK: *A. incana, A. cortada, A. rubra*, and *A. viridis*, and hybrid populations have been identified (Figure S19) (Banaev and Bažant 2007; Baxter and McAllister 2024; Gerchen et al. 2026). The suspected hybrid individuals lie within an area where multiple *Alnus* species and known hybrid distributions overlap (Figure S19). But this overlap covers much of the UK, including populations with no evidence of hybridisation in our dataset. Species distributions alone therefore cannot confirm that these are natural hybrids, and as contamination is another likely cause, we did not investigate these samples further.

We identified two triploids, all other individuals being diploid. These may have arisen through unreduced gamete formation or hybridisation of diploid and tetraploid *Alnus* species, as has been found in European mixed-ploidy contact zones (Banaev and Bažant 2007; Šmíd et al. 2020). Darlington & Wylie (1955) reported most *Alnus* species as diploid, with no triploids, but with some tetraploids identified for *A. glutinosa*. Both diploids and triploids (3x = 42) were reported for *A. cordata*, *A. hirsuta*, *A. orientalis*, *A. japonica*, and *A. subcordata* (Darlington and Wylie 1955). Triploidy in our dataset is therefore biologically plausible, with various polyploid *Alnus* cytotypes previously reported (Banaev and Bažant 2007; Šmíd et al. 2020; Darlington and Wylie 1955).

Three main lineages are recognised in the *Alnus glutinosa* aggregate: diploid *A. glutinosa*, and the newly described tetraploid *A. rohlenae* from the Balkans and tetraploid *A. lusitanica* from the Iberian Peninsula, both of which were previously thought to be diploid *A. glutinosa* (Vít et al. 2017; Gomes Marques et al. 2022; Martín et al. 2024). Hybridisation has been found in Europe between *A. glutinosa* and various other species: with *A. cordata* (2x) in Italy (Villani et al. 2021; Pollegioni et al. 2025), with *A. incana* (2x) in Belgium (Vander Mijnsbrugge 2015), with *A. rohlenae* (4x) in Greece (Vít et al. 2017), and *A. lusitanic*a (4x) in Spain (Banaev and Bažant 2007; Šmíd et al. 2020; Gerchen et al. 2026). These Spanish hybrids were adult triploid trees, but a study published just two years earlier, which sampled populations from the same mixed ploidy contact zone in the river basin, did not identify hybrids (Martín et al. 2024). Given this complexity within the genus, with hybridisation and gene flow across and between lineages of various ploidy, further investigation is needed to understand the origin of the triploids identified.

We found a single individual with evidence for trisomy. Prior to whole genome sequencing, trisomy would have been very hard to detect. This is the first evidence we are aware of for its occurrence in this species. Trisomy is unlikely to be a consequence of hybridisation and is most likely due to errors in meiosis. Whether this seedling would have survived to adulthood we do not know.

### UK *Alnus glutinosa* population structure

We found little population structure in the majority of UK alder populations (Figure 2), apart from that caused by two northern populations (924731 and 854432). Low differentiation is expected in a wind-pollinated species with water- and wind-dispersed seeds, and has been shown in previous alder studies, including microsatellite analysis of Northern Irish populations (Beatty et al. 2015) and genotyping of Irish plus-trees (Huang et al. 2026). In the UK, we found evidence that genetic diversity declined with increasing latitude whilst population differentiation increased, consistent with genetic drift and founder effects as a result of northwards colonisation following the last glacial maximum. The separation of population 924731 alongside excess of heterozygosity could reflect gene flow from unsampled, genetically distinct neighbouring populations. The genetic status of population 854432 remains unclear, given its separation in PCA space and outlying pairwise F_ST_ estimates which may be influenced by sample size.

### Environmental variables with greatest genomic signal

Across 7.38 million genome-wide SNPs, LFMMs identified 1,364 significant SNPs of which 1,269 were associated with the average number of snow days. Whilst it is tempting to conclude that snow is therefore a major driver of local adaptation in UK *A. glutinosa* populations, this variable showed high levels of genomic inflation (Figure S11), and allele frequency clines for the significant alleles were strongly affected by outlier populations (Figure S13). Thus, the number of loci crossing the FDR threshold for this variable is not a reliable indication of its importance to local adaptation. Significant SNPs for other environmental variables also contained outlier populations which could be causing false positives (Figure S13).

To overcome this burden and accurately assess the contribution of environmental variables on adaptation, we utilised an approach developed by Carleial et al <u>(2025)</u> (Carleial et al. 2025), where all 7.38 million genome-wide loci were used to generate a covariance matrix of SNP-effect sizes from which the total, shared and unique genomic signal contributed by each variable could be estimated. Adopting this method, the signal attributed to snow days was moderate, as effect sizes co-varied with other environmental variables, leading to a relatively low unique genomic signal. Mean temperature of the wettest quarter, annual mean frost days and precipitation of the driest month explained the greatest whole-genome variance in SNP effect sizes while also showing a high number of single-loci significant associations, and were therefore supported by both approaches. These variables represent thermal and hydrological extremes, consistent with the ecology of alder in riparian habitats, where adaptation to periods of drought, flooding and temperature extremes is likely important. Restoration programmes and climate-informed management should therefore prioritise sourcing seeds from locations matching the temperature and hydrological profile of the planting site to link adapted allele frequencies to adaptive local climates, thereby promoting a ‘right tree, right place’ approach.

Our results also emphasise the importance of developing appropriate null models for genetic association studies. Genomic signals explained by easting and isothermality were not significantly higher than that explained by a dummy variable despite signals being largely independent from other variables Yet, for isothermality, many SNPs crossed the FDR significant threshold in the single-locus GEA. These results add to an increasing body of literature showing that GEAs can produce many false positives despite stringent significant thresholds and carefully controlling for confounding factors such as population structure and kinship (Lotterhos 2023; Booker et al. 2024; Lazic et al. 2024; Carleial et al. 2025). Like previous studies, we suggest that future work should move away from focusing on arbitrary ‘significant’ SNPs, and instead make use of genome-wide approaches to better characterise the genetic architecture of important traits.

### Predicting genetic change required for adaptation

While estimating the genetic change required for present-day populations to match future climates (i.e. genomic offsets), we found a significant relationship between genomic offset and latitude using both the *LEA* and *AlleleShift* methods. Estimates from *LEA* were significantly associated with climate dissimilarity, but latitude was a much better predictor. This suggests our estimates of climate dissimilarity may have failed to include meaningful environmental variables which are indirectly captured by latitudinal gradients (phenology, growing season length, soil characteristics). Future GEAs should include soil characteristics such as moisture content as these variables are worth considering when selecting seed stock for resilient planting (Hrivnák et al. 2023).

The higher genomic offset estimates of northern populations to current and future climates suggests they are less adapted to local conditions, and may be genetically disadvantaged against future change. This may have arisen due to genetic drift during northwards recolonisation, or due to selection of alleles that confer an advantage in colder environments but that are maladaptive to future, warmer, climates. To mitigate climate-driven population declines, seed for planting in northern locations could be sourced from southern sites, thereby promoting gene flow of beneficial alleles to genetically vulnerable populations. As stated above, variables related to temperature and precipitation should be the main focus when choosing appropriate sites. This assisted gene flow approach should also increase the genetic diversity of northern populations, enhancing resilience to future conditions.

Future work includes implementing alternative GEA approaches to validate environmental variables involved in local adaptation (Ahrens et al. 2021; Capblancq et al. 2018; Lazic et al. 2024; Vanhove et al. 2021). Additionally, the establishment of common garden experiments, such as those completed by Kort et al (2014), would help strengthen the link between genotype, phenotype and the environment (De Kort et al. 2014; Villoutreix et al. 2025; Wu et al. 2026), as well as validate genomic offset approaches (Rellstab and Keller 2025; Carleial et al. 2025). Further analyses could identify seed sources predicted to be best adapted to the future climates of target planting sites, thereby informing climate-aware seed sourcing by helping select planting material with lower predicted genomic offset under future climates (Grabowski et al. 2025).

## Acknowledgements

We thank Gabriele Nocchi for providing assistance and resources for the analyses in this paper. Funding was provided by the UK Department for Environment, Food and Rural Affairs via the Centre for Forest Protection. This research utilised Queen Mary’s Apocrita HPC facility, supported by QMUL Research-IT (http://doi.org/10.5281/zenodo.438045).

## Data Accessibility

DNA sequences: NCBI BioProject accession PRJNA825844

Reference Genome and Annotation: Darwin Tree of Life (DToL) reference genome of *Alnus glutinosa*; Assembly version GCA_958979055.1, available from https://www.ebi.ac.uk/ena/browser/view/GCA_958979055 (Darwin Tree of Life Project, n.d.).

Climate data: Bio1-Bio19 climatologies from CHELSA V2.1 database available from https://www.chelsa-climate.org/models/chelsa (Karger et al. 2017); MetOffice HadUK-Grid datasets available at https://www.metoffice.gov.uk/research/climate/maps-and-data/data/haduk-grid/haduk-grid (Met Office et al. 2024); terrain characteristics available from https://biogeo.ucdavis.edu/data/diva/alt/GBR_alt.zip ; MetOffice UK Climate Projections (UKPC) database (Met Office) available at https://ukclimateprojections-ui.metoffice.gov.uk/ui/home; MetOffice climate data portal (Met Office, 2023) available at https://climatedataportal.metoffice.gov.uk/datasets/TheMetOffice::annual-count-of-frost-days-projections-12km/about.

Code: Will be made available upon publication

## Author Contributions

The project was designed by R.B and M.C. Seed collection and seed bank access was provided by T.C. Germination and growing was performed by The Jodrell Glasshouse Team, Kew and M.C. Genomic analysis, manuscript writing and figure creation was performed by P.S. Genomic analysis was supported by R.C. Manuscript writing was supported by R.B and R.C.

## References

Ahrens, Collin W., Rebecca Jordan, Jason Bragg, et al. 2021. “Regarding the F-Word: The Effects of Data Filtering on Inferred Genotype-Environment Associations.” Molecular Ecology Resources 21 (5): 1460–1474.

Alexander, David H., and Kenneth Lange. 2011. “Enhancements to the ADMIXTURE Algorithm for Individual Ancestry Estimation.” BMC Bioinformatics 12 (1): 246.

Andrews, S. 2010. “FastQC: A Quality Control Tool for High Throughput Sequence Data.” https://www.bioinformatics.babraham.ac.uk/projects/fastqc/.

Banaev, V., and V. Bažant. 2007. “Study of Natural Hybridization between Alnus Incana (L.) Moench. and Alnus Glutinosa (L.) Gaertn.” Journal of Forest Science 53 (2): 66–73.

Bass, A., Storey, J., Chirico, M. & Kryukov, I. 2023. Qvalue: R Package to Estimate Q-Values and False Discovery Rate Quantities. Github. https://github.com/StoreyLab/qvalue.

Baxter, T. & McAllister, H.A. *Trees and Shrubs Online* (treesandshrubsonline.Org/articles/Alnus/Alnus-Glutinosa/). 2024. “Alnus Glutinosa (L.) Gaertn.” https://www.treesandshrubsonline.org/articles/alnus/alnus-glutinosa/.

Beatty, Gemma E., W. Ian Montgomery, David G. Tosh, and Jim Provan. 2015. “Genetic Provenance and Best Practice Woodland Management: A Case Study in Native Alder (Alnus Glutinosa).” Tree Genetics & Genomes 11 (5). 10.1007/s11295-015-0919-1.

Beheregaray, Luciano B., Sarah W. Fitzpatrick, Andrew R. Whiteley, Jill Anderson, and Paul Sunnucks. 2026. “Genetic Rescue: Latest Advances and Applications.” Evolutionary Applications 19 (3): e70225.

Bolger, Anthony M., Marc Lohse, and Bjoern Usadel. 2014. “Trimmomatic: A Flexible Trimmer for Illumina Sequence Data.” *Bioinformatics (Oxford*, England) 30 (15): 2114–2120.

Booker, Tom R., Sam Yeaman, James R. Whiting, and Michael C. Whitlock. 2024. “The WZA: A Window-Based Method for Characterizing Genotype-Environment Associations.” Molecular Ecology Resources 24 (2): e13768.

Browning, Brian L., Ying Zhou, and Sharon R. Browning. 2018. “A One-Penny Imputed Genome from next-Generation Reference Panels.” American Journal of Human Genetics 103 (3): 338–348.

Camacho, Christiam, George Coulouris, Vahram Avagyan, et al. 2009. “BLAST+: Architecture and Applications.” BMC Bioinformatics 10 (1): 421.

Capblancq, Thibaut, Keurcien Luu, Michael G. B. Blum, and Eric Bazin. 2018. “Evaluation of Redundancy Analysis to Identify Signatures of Local Adaptation.” Molecular Ecology Resources 18 (6): 1223–1233.

Carleial, Rômulo, Michael Charters, Domen Finzgar, et al. 2025. “Genetic Basis of Traits and Local Adaptation in UK Silver Birch.” In bioRxiv. 10.1101/2025.07.04.662749.

Catchen, Julian, Paul A. Hohenlohe, Susan Bassham, Angel Amores, and William A. Cresko. 2013. “Stacks: An Analysis Tool Set for Population Genomics.” Molecular Ecology 22 (11): 3124–3140.

Caye, Kevin, Basile Jumentier, Johanna Lepeule, and Olivier François. 2019. “LFMM 2: Fast and Accurate Inference of Gene-Environment Associations in Genome-Wide Studies.” Molecular Biology and Evolution 36 (4): 852–860.

Chang, Christopher C., Carson C. Chow, Laurent Cam Tellier, Shashaank Vattikuti, Shaun M. Purcell, and James J. Lee. 2015. “Second-Generation PLINK: Rising to the Challenge of Larger and Richer Datasets.” GigaScience 4 (1): 7.

Chapman, Ted. 2024. “UK National Tree Seed Database and Portal.” RBG Kew. https://www.kew.org/science/our-science/projects/uk-national-tree-seed-database-and-portal.

Chen, Yuxin, Yongsheng Chen, Chunmei Shi, et al. 2018. “SOAPnuke: A MapReduce Acceleration-Supported Software for Integrated Quality Control and Preprocessing of High-Throughput Sequencing Data.” GigaScience 7 (1): 1–6.

Claessens, H., A. Oosterbaan, P. Savill, and J. Rondeux. 2010. “A Review of the Characteristics of Black Alder (Alnus Glutinosa (L.) Gaertn.) and Their Implications for Silvicultural Practices.” *Forestry (London*, England) 83 (2): 163–175.

Committee on Climate Change. 2019. Net Zero: The UK’s Contribution to Stopping Global Warming. Committee on Climate Change. https://www.theccc.org.uk/wp-content/uploads/2019/05/Net-Zero-The-UKs-contribution-to-stopping-global-warming.pdf.

Cox, K., A. Vanden Broeck, H. Van Calster, and J. Mergeay. 2011. “Temperature-Related Natural Selection in a Wind-Pollinated Tree across Regional and Continental Scales: SELECTIVE LOCI VS. CLIMATE IN ALNUS GLUTINOSA.” Molecular Ecology 20 (13): 2724–2738.

Danecek, Petr, Adam Auton, Goncalo Abecasis, et al. 2011. “The Variant Call Format and VCFtools.” Bioinformatics (Oxford, England) 27 (15): 2156–2158.

Darlington, C. D., and A. P. Wylie. 1955. Chromosome Atlas of Flowering Plants. George Allen & Unwin Ltd.

Darwin Tree of Life Project. “Assembly: GCA_958979055.1.” European Nucleotide Archive. Accessed June 2, 2026. https://www.ebi.ac.uk/ena/browser/view/GCA_958979055.

De Kort, Hanne, Katrien Vandepitte, Hans Henrik Bruun, Déborah Closset-Kopp, Olivier Honnay, and Joachim Mergeay. 2014. “Landscape Genomics and a Common Garden Trial Reveal Adaptive Differentiation to Temperature across Europe in the Tree Species Alnus Glutinosa.” Molecular Ecology 23 (19): 4709–4721.

DePristo, Mark A., Eric Banks, Ryan Poplin, et al. 2011. “A Framework for Variation Discovery and Genotyping Using next-Generation DNA Sequencing Data.” Nature Genetics 43 (5): 491–498.

Edinburgh Genetics. n.d. “EgSEQ Enzymatic Library Prep.” Accessed June 2, 2026. https://www.eggenetics.com/library-preparation/egseq-enzymatic-library-prep.

Ewels, Philip, Måns Magnusson, Sverker Lundin, and Max Käller. 2016. “MultiQC: Summarize Analysis Results for Multiple Tools and Samples in a Single Report.” Bioinformatics (Oxford, England) 32 (19): 3047–3048.

Ferguson, T. P., and G. Bond. 1953. “Observations on the Formation and Function of the Root Nodules of Alnus Glutinosa (L.) Gaertn.” Annals of Botany 17 (1): 175–188.

Forget, Guillaume, Clélia Carreau, Didier Le Coeur, and Ivan Bernez. 2013. “Ecological Restoration of Headwaters in a Rural Landscape (Normandy, France): A Passive Approach Taking Hedge Networks into Account for Riparian Tree Recruitment.” Restoration Ecology 21 (1): 96–104.

Frichot, Eric, Olivier Francois, and Clement Gain. 2026. “Lea: An R Package for Landscape and Ecological Association Studies.” Preprint. https://bioc.r-universe.dev/LEA/LEA.pdf.

Frichot, Eric, François Mathieu, Théo Trouillon, Guillaume Bouchard, and Olivier François. 2014. “Fast and Efficient Estimation of Individual Ancestry Coefficients.” Genetics 196 (4): 973–983.

Gain, Clément, Bénédicte Rhoné, Philippe Cubry, et al. 2023. “A Quantitative Theory for Genomic Offset Statistics.” Molecular Biology and Evolution 40 (6): msad140.

García-Alcalde, Fernando, Konstantin Okonechnikov, José Carbonell, et al. 2012. “Qualimap: Evaluating next-Generation Sequencing Alignment Data.” Bioinformatics (Oxford, England) 28 (20): 2678–2679.

Gerchen, Jörn F., Bohumil Mandák, Milena Melnyk, and Filip Kolář. 2026. “Centromeric Variation Is Shared across Ploidy Barriers in *Alnus Glutinosa* Agg.” In bioRxiv. BioRxiv, April 23. 10.64898/2026.04.21.719804.

Gogarten, Stephanie M., Tushar Bhangale, Matthew P. Conomos, et al. 2012. “GWASTools: An R/Bioconductor Package for Quality Control and Analysis of Genome-Wide Association Studies.” Bioinformatics (Oxford, England) 28 (24): 3329–3331.

Gomes Marques, Inês, Carla Faria, Sofia Isabel Rodrigues Conceição, et al. 2022. “Germination and Seed Traits in Common Alder ( *Alnus* Spp.): The Potential Contribution of Rear edge Populations to Ecological Restoration Success.” Restoration Ecology 30 (3). 10.1111/rec.13517.

Grabowski, Paul P., Chloee M. McLaughlin, Joanna Rifkin, et al. 2025. “Leveraging Species-Wide Variation and Patterns of Adaptation to Inform Pecan Crop Improvement Efforts.” In bioRxiv. BioRxiv, December 22. 10.64898/2025.12.19.695279.

Hijmans, Robert J. 2025. “Geographic Data Analysis and Modeling [R Package Raster Version 3.6-32].” Comprehensive R Archive Network (CRAN), March 28. https://cran.r-project.org/web/packages/raster/index.html.

Hrivnák, Matúš, Diana Krajmerová, Richard Hrivnák, et al. 2023. “Interplay between Tree Genetic Variation, Plant Community Composition and Environment in Forest Communities Dominated by Black Alder (Alnus Glutinosa (L.) Gaertn.).” Perspectives in Plant Ecology, Evolution and Systematics 60 (125748): 125748.

Huang, Jie, Stephen Byrne, Oliver Sheridan, et al. 2026. “Genetic Diversity Assessment and Genomic Prediction in Alder (Alnus Glutinosa) to Support Seed Orchard Improvement and Sustainable Forestry.” Discover Plants 3 (1). 10.1007/s44372-026-00862-z.

Jombart, Thibaut. 2008. “Adegenet: A R Package for the Multivariate Analysis of Genetic Markers.” Bioinformatics (Oxford, England) 24 (11): 1403–1405.

Juliusdottir, Thorhildur. 2023. “Topr: An R Package for Viewing and Annotating Genetic Association Results.” BMC Bioinformatics 24 (1): 268.

Karger, Dirk Nikolaus, Olaf Conrad, Jürgen Böhner, et al. 2017. “Climatologies at High Resolution for the Earth’s Land Surface Areas.” Scientific Data 4 (1): 170122.

Kassambara, Alboukadel. 2022. Ggcorrplot: Visualization of a Correlation Matrix Using ggplot2. Github. https://github.com/kassambara/ggcorrplot.

Kawecki, Tadeusz J., and Dieter Ebert. 2004. “Conceptual Issues in Local Adaptation.” Ecology Letters 7 (12): 1225–1241.

Kindt, Roeland. 2021a. “AlleleShift: An R Package to Predict and Visualize Population-Level Changes in Allele Frequencies in Response to Climate Change.” PeerJ 9 (e11534): e11534.

Kindt, Roeland. 2021b. “AlleleShift: Predict and Visualize Population-Level Changes in Allele Frequencies in Response to Climate Change.” In CRAN: Contributed Packages. The R Foundation, January 26. 10.32614/cran.package.alleleshift.

Knaus, Brian J., and Niklaus J. Grünwald. 2017. “Vcfr: A Package to Manipulate and Visualize Variant Call Format Data in R.” Molecular Ecology Resources 17 (1): 44–53.

Kuhn, Max, Simon Jackson, and Jorge Cimentada. 2016. “Corrr: Correlations in R.” In *CRAN: Contributed Packages*. The R Foundation, July 8. 10.32614/cran.package.corrr.

Laetsch, Dominik, and Mark Blaxter. 2017. “BlobTools: Interrogation of Genome Assemblies.” In F1000Res. 10.12688/f1000research.12232.1.

Lasky, Jesse R., Emily B. Josephs, and Geoffrey P. Morris. 2023. “Genotype-Environment Associations to Reveal the Molecular Basis of Environmental Adaptation.” The Plant Cell 35 (1): 125–138.

Lazic, Desanka, Cornelia Geßner, Katharina J. Liepe, et al. 2024. “Genomic Variation of European Beech Reveals Signals of Local Adaptation despite High Levels of Phenotypic Plasticity.” Nature Communications 15 (1): 8553.

Leroy, Boris, Christine N. Meynard, Céline Bellard, and Franck Courchamp. 2016. “Virtualspecies, an R Package to Generate Virtual Species Distributions.” Ecography 39 (6): 599–607.

Liang, Yi-Ye, Hui Liu, Qiong-Qiong Lin, et al. 2025. “Pan-Genome Analysis Reveals Local Adaptation to Climate Driven by Introgression in Oak Species.” Molecular Biology and Evolution 42 (5): msaf088.

Li, Heng, and Richard Durbin. 2009. “Fast and Accurate Short Read Alignment with Burrows-Wheeler Transform.” Bioinformatics (Oxford, England) 25 (14): 1754–1760.

Li, Heng, Bob Handsaker, Alec Wysoker, et al. 2009. “The Sequence Alignment/Map Format and SAMtools.” *Bioinformatics (Oxford*, England) 25 (16): 2078–2079.

Long, Zhiqin, Yupeng Sang, Jiajun Feng, et al. 2025. “Evolutionary Genomics Unravels the Responses and Adaptation to Climate Change in a Key Alpine Forest Tree Species.” Molecular Biology and Evolution 42 (7): msaf116.

Lotterhos, Katie E. 2023. “The Paradox of Adaptive Trait Clines with Nonclinal Patterns in the Underlying Genes.” Proceedings of the National Academy of Sciences of the United States of America 120 (12): e2220313120.

Martín, M. Angela, R. Moreno, J. V. Die, et al. 2024. “Distribution, Diversity and Genetic Structure of Alders (Alnus Lusitanica and A. Glutinosa) in Spain.” Forest Ecology and Management 562 (121922): Not Available.

McVean, D. N. 1953. “Alnus Glutinosa (L.) Gaertn.” The Journal of Ecology 41 (2): 447– 466.

Met Office. 2023. “Annual Count of Frost Days - Projections (12km).” Met Office Climate Data Portal. https://climatedataportal.metoffice.gov.uk/datasets/TheMetOffice::annual-count-of-frost-days-projections-12km/about.

Met Office. “UK Climate Projections (UKCP18).” Accessed June 3, 2026. https://www.metoffice.gov.uk/research/approach/collaboration/ukcp.

Met Office, Dan Hollis, Emily Carlisle, Michael Kendon, Stephen Packman, and Amy Doherty. 2024. “HadUK-Grid Gridded Climate Observations on a 1km Grid over the UK, v1.3.0.ceda (1836-2023).” NERC EDS Centre for Environmental Data Analysis, July 18. 10.5285/B963EAD70580451AA7455782224479D5.

Nakamura, Futoshi. 2022. “Riparian Forests and Climate Change: Interactive Zone of Green and Blue Infrastructure.” In Ecological Research Monographs. Springer Singapore.

Patel, Ravi K., and Mukesh Jain. 2012. “NGS QC Toolkit: A Toolkit for Quality Control of next Generation Sequencing Data.” PloS One 7 (2): e30619.

Pollegioni, Paola, Alexis Marchesini, Muriel Gaudet, et al. 2025. “What Shapes the Genetic Diversity of the Alnus Cordata Species across Its Italian Native Range? Informing Conservation Strategies.” Ecology and Evolution 15 (9): e72018.

Poplin, Ryan, Valentin Ruano-Rubio, Mark DePristo, et al. 2017. “Scaling Accurate Genetic Variant Discovery to Tens of Thousands of Samples.” In bioRxiv. BioRxiv. 10.1101/201178.

Raj, Anil, Matthew Stephens, and Jonathan K. Pritchard. 2014. “fastSTRUCTURE: Variational Inference of Population Structure in Large SNP Data Sets.” Genetics 197 (2): 573–589.

R Core Team. 2026. R: A Language and Environment for Statistical Computing. R Foundation for Statistical Computing.

Rellstab, Christian, Felix Gugerli, Andrew J. Eckert, Angela M. Hancock, and Rolf Holderegger. 2015. “A Practical Guide to Environmental Association Analysis in Landscape Genomics.” Molecular Ecology 24 (17): 4348–4370.

Rellstab, Christian, and Stephen R. Keller. 2025. “Can We Use Genomic Data to Predict Maladaptation to Environmental Change?” Molecular Ecology Resources 25 (4): e14059.

Sang, Yupeng, Zhiqin Long, Xuming Dan, et al. 2022. “Genomic Insights into Local Adaptation and Future Climate-Induced Vulnerability of a Keystone Forest Tree in East Asia.” Nature Communications 13 (1): 6541.

Savolainen, Outi, Martin Lascoux, and Juha Merilä. 2013. “Ecological Genomics of Local Adaptation.” Nature Reviews. Genetics 14 (11): 807–820.

Šmíd, Jan, Jan Douda, Karol Krak, and Bohumil Mandák. 2020. “Analyses of Hybrid Viability across a Hybrid Zone between Two Alnus Species Using Microsatellites and cpDNA Markers.” Genes 11 (7): E770.

Vacek, Zdeněk, Stanislav Vacek, Jan Cukor, et al. 2022. “Dendrochronological Data from Twelve Countries Proved Definite Growth Response of Black Alder (*Alnus Glutinosa* [L.] Gaertn.) to Climate Courses across Its Distribution Range.” Central European Forestry Journal 68 (3): 139–153.

Vaht, Riina, Margus Pensa, Mait Sepp, Aarne Luud, Helen Karu, and Tiina Elvisto. 2010. “Assessment of Vegetation Performance on Semicoke Dumps of Kohtla-Järve Oil Shale Industry, Estonia; Pp. 3–18.” Estonian Journal of Ecology 59 (1): 3–18.

Van Der Auwera. GA, O’connorc B. D. 2020. Genomics in the Cloud: Using Docker, GATK, and WDL in Terra (1st Edition). O’Reilly Media.

Vander Mijnsbrugge, Kristine. 2015. “Morphological Dissection of Leaf, Bud and Infructescence Traits of the Interfertile Native A. Glutinosa and Non-Native A. Incana in Flanders (northern Part of Belgium).” Trees (Berlin, Germany: West) 29 (6): 1661– 1672.

Vanhove, Mathieu, Francisco Pina-Martins, Ana Cristina Coelho, et al. 2021. “Using Gradient Forest to Predict Climate Response and Adaptation in Cork Oak.” Journal of Evolutionary Biology 34 (6): 910–923.

Villani, Fiorella, Simone Castellana, Isacco Beritognolo, et al. 2021. “Genetic Variability of Alnus Cordata (Loisel.) Duby Populations and Introgressive Hybridization with A. Glutinosa (L.) Gaertn. In Southern Italy: Implication for Conservation and Management of Genetic Resources.” Forests 12 (6): 655.

Villanueva, Randle Aaron M., and Zhuo Job Chen. 2019. “ggplot2: Elegant Graphics for Data Analysis (2nd Ed.).” Measurement: Interdisciplinary Research and Perspectives 17 (3): 160–167.

Villoutreix, Romain, Nathalie Faure, Cédric Glorieux, and Fabrice Roux. 2025. “Predicting Fitness in Future Climates: Insights from Temporally Replicated Field Experiments in Arabidopsis Thaliana.” Evolution Letters 9 (4): 392–407.

Vít, Petr, Jan Douda, Karol Krak, Alena Havrdová, and Bohumil Mandák. 2017. “Two New Polyploid Species Closely Related to *Alnus Glutinosa* in Europe and North Africa – An Analysis Based on Morphometry, Karyology, Flow Cytometry and Microsatellites.” Taxon 66 (3): 567–583.

Wei, Taiyun, and Viliam Simko. 2010. “Corrplot: Visualization of a Correlation Matrix.” In *CRAN: Contributed Packages*. The R Foundation, April 10. 10.32614/cran.package.corrplot.

Whitlock, Michael C. 2015. “Modern Approaches to Local Adaptation.” The American Naturalist 186 Suppl 1 (S1): S1–4.

Wu, Xing, Tatiana Bellagio, Yunru Peng, et al. 2026. “Rapid Adaptation and Extinction in Synchronized Outdoor Evolution Experiments of Arabidopsis.” Science (New York, N.Y.) 391 (6792): eadz0777.

Yin, Lilin, Haohao Zhang, Zhenshuang Tang, et al. 2021. “RMVP: A Memory-Efficient, Visualization-Enhanced, and Parallel-Accelerated Tool for Genome-Wide Association Study.” *Genomics*, Proteomics & Bioinformatics 19 (4): 619–628.

Zhang, Jiangli, Andrea Ghirardo, Antonella Gori, et al. 2020. “Improving Air Quality by Nitric Oxide Consumption of Climate-Resilient Trees Suitable for Urban Greening.” Frontiers in Plant Science 11 (September): 549913.

