## Supplement for "Genomic signals of local adaptation in UK black alder populations (*Alnus glutinosa*)"

1 – Royal Botanic Gardens Kew, Richmond, Surrey, TW9 3AE, UK

2 – Queen Mary University of London, Mile End Road, London, E1 4NS

| Population ID | Latitude (degrees) | Longitude (degrees) | Number of samples sequences | Number of diploid samples in GEA | Estimated Population Size by UKNTSP |
|---|---|---|---|---|---|
| 952006 | 50.50 | -4.03 | 3 | 3 | 50 |
| 904120 | 50.68 | -3.69 | 5 | 5 | >80 |
| 815549 | 50.76 | -2.92 | 10 | 10 | 100 |
| 870104 | 50.92 | -4.51 | 4 | 4 | >50 |
| 758725 | 51.06 | -0.10 | 13 | 13 | 300 |
| 763064 | 51.06 | -0.099 | 8 | 8 | 300 |
| 770857 | 51.41 | -1.52 | 10 | 10 | 40 |
| 954583 | 51.52 | -3.17 | 14 | 14 | 100 |
| 905150 | 51.74 | -3.17 | 15 | 14 | 100 |
| 855288 | 51.88 | -3.02 | 3 | 3 | >1000 |
| 905172 | 51.94 | -3.04 | 13 | 13 | 40 |
| 1013696 | 51.94 | -3.03 | 8 | 8 | >100 |
| 925026 | 52.00 | -0.47 | 11 | 11 | 32 |
| 899419 | 52.17 | 0.82 | 14 | 14 | 30 |
| 850102 | 52.37 | -1.43 | 4 | 4 | 10 |
| 850098 | 52.37 | -1.43 | 7 | 7 | 15 |
| 850113 | 52.37 | -1.43 | 1 | 1 | 5 |
| 899327 | 52.88 | -3.11 | 6 | 6 | 25 |
| 865212 | 53.32 | -1.65 | 11 | 10 | >60 |
| 954538 | 54.16 | -7.45 | 10 | 7 | 30 |
| 849377 | 54.26 | -1.81 | 4 | 4 | N.A |
| 1012482 | 54.33 | -2.74 | 2 | 2 | 10 |
| 1012563 | 54.37 | -2.90 | 1 | 1 | 10 |
| 954527 | 54.44 | -7.72 | 20 | 20 | 30 |
| 951881 | 54.44 | -7.74 | 2 | 2 | 10 |
| 1012677 | 54.44 | -3.00 | 4 | 4 | N.A |
| 851947 | 54.52 | -0.84 | 6 | 6 | 20 |
| 861395 | 55.05 | -4.55 | 5 | 5 | >100 |

| | | | | | |
|---|---|---|---|---|---|
| 899305 | 55.08 | -4.45 | 11 | 11 | >80 |
| 956705 | 55.36 | -2.65 | 10 | 10 | 30 |
| 954365 | 55.53 | -2.78 | 2 | 2 | 30 |
| 899338 | 55.65 | -3.77 | 1 | 1 | >50 |
| 954435 | 55.73 | -3.18 | 7 | 7 | >20 |
| 911452 | 55.84 | -5.24 | 1 | 1 | >30 |
| 899279 | 55.99 | -3.81 | 14 | 7 | 50 |
| 849953 | 55.99 | -5.20 | 1 | 1 | 300 |
| 912426 | 56.24 | -3.23 | 3 | 3 | >100 |
| 870115 | 56.27 | -3.23 | 1 | 1 | 9 |
| 895710 | 56.42 | -4.46 | 13 | 13 | N.A |
| 854432 | 57.17 | -4.84 | 26 | 26 | 60 |
| 924720 | 57.21 | -3.72 | 4 | 3 | 20 |
| 925473 | 57.22 | -3.69 | 1 | 1 | 8-10 |
| 926768 | 57.37 | -4.39 | 1 | 1 | 12 |
| 924731 | 57.55 | -4.49 | 6 | 6 | >100 |

**Table S1 – Details for each of the 44 UK A*lnus glutinosa* accessions genotyped.** Columns detail coordinates, number of genotyped samples, number of samples following polyploid and hybrid removal, estimated population size as provided by the UKNTSP during seed collection, and sample numbers in plots.

| | *Predictors* | *Estimates* | *CI* | *F-Statistic* | *pvalue* |
|---|---|---|---|---|---|
| **P** $R^2$ / $R^2$ adjusted : 0.143 / 0.122 | (Intercept) | 0.68 | 0.55 – 0.81 | - | - |
| | LAT | <0.001 | 0.00 – 0.01 | 7 | **0.011** |
| **Obs_Het** $R^2$ / $R^2$ adjusted : 0.001 / -0.022 | (Intercept) | 0.25 | 0.17 – 0.33 | - | - |
| | LAT | <0.001 | -0.00 – 0.00 | 0.06 | 0.806 |
| **Exp_Het** $R^2$ / $R^2$ adjusted : 0.161 / 0.141 | (Intercept) | 0.64 | 0.33 – 0.95 | - | - |
| | LAT | -0.01 | -0.01 – -0.00 | 8.05 | **0.007** |
| **Pi** $R^2$ / $R^2$ adjusted : 0.022 / -0.001 | (Intercept) | 0.28 | 0.20 – 0.36 | - | - |
| | LAT | <0.001 | -0.00 – 0.00 | 0.95 | 0.337 |
| **Fis** $R^2$ / $R^2$ adjusted : 0.084 / 0.063 | (Intercept) | 0.09 | 0.00 – 0.18 | - | - |
| | LAT | <0.001 | -0.00 – 0.00 | 3.88 | 0.056 |

**Table S2 – Relationship between population latitude and genetic diversity metrics.** STACKS populations was used to estimate measures of genetic diversity across 304 individuals from 44 populations. These were then regressed against population latitude (Figure S7). Raw values are shown in Figure S6.

| Environmental Variable | Description |
|---|---|
| Precipitation of the Driest Month | Minimum monthly precipitation total (kg $m^{-2}$ $month^{-1}$) |
| Isothermality | 100 × (diurnal temperature range ÷ daily temperature range); compares day–night variability to annual temperature range (°C) |
| Mean Temperature of the Wettest Quarter | Average temperature of the wettest four months of the year (°C) |
| Mean Temperature of the Driest Quarter | Average temperature of the driest four months of the year (°C) |
| Slope | Rate of elevation change between each focal cell and its eight surrounding cells |
| TPI (Topographic Position Index) | Topographic Position Index - difference between the value of a cell and the mean value of its 8 surrounding cells |
| Annual Mean Frost Days | Count of days when the grass minimum temperature is below 0ºC (days) |
| Average Wind Speed | Average of hourly mean wind speed at a height of 10 m above ground level (knots) |
| Average Snow Days | Count of days with greater than 50% of the ground covered by snow at 0900 UTC (days) |
| Northing | Degree to which a slope faces north or south. Calculated as the cosine of the aspect terrain variable (°) |
| Easting | Degree to which a slope faces east or west. Calculated as the sine of the aspect terrain variable (°) |

**Table S3 – Minimally collinear environmental variables used in the GEA**. List includes climate and terrain variables remaining following pruning of collinear variables as described in methods and presented in Figure S8.

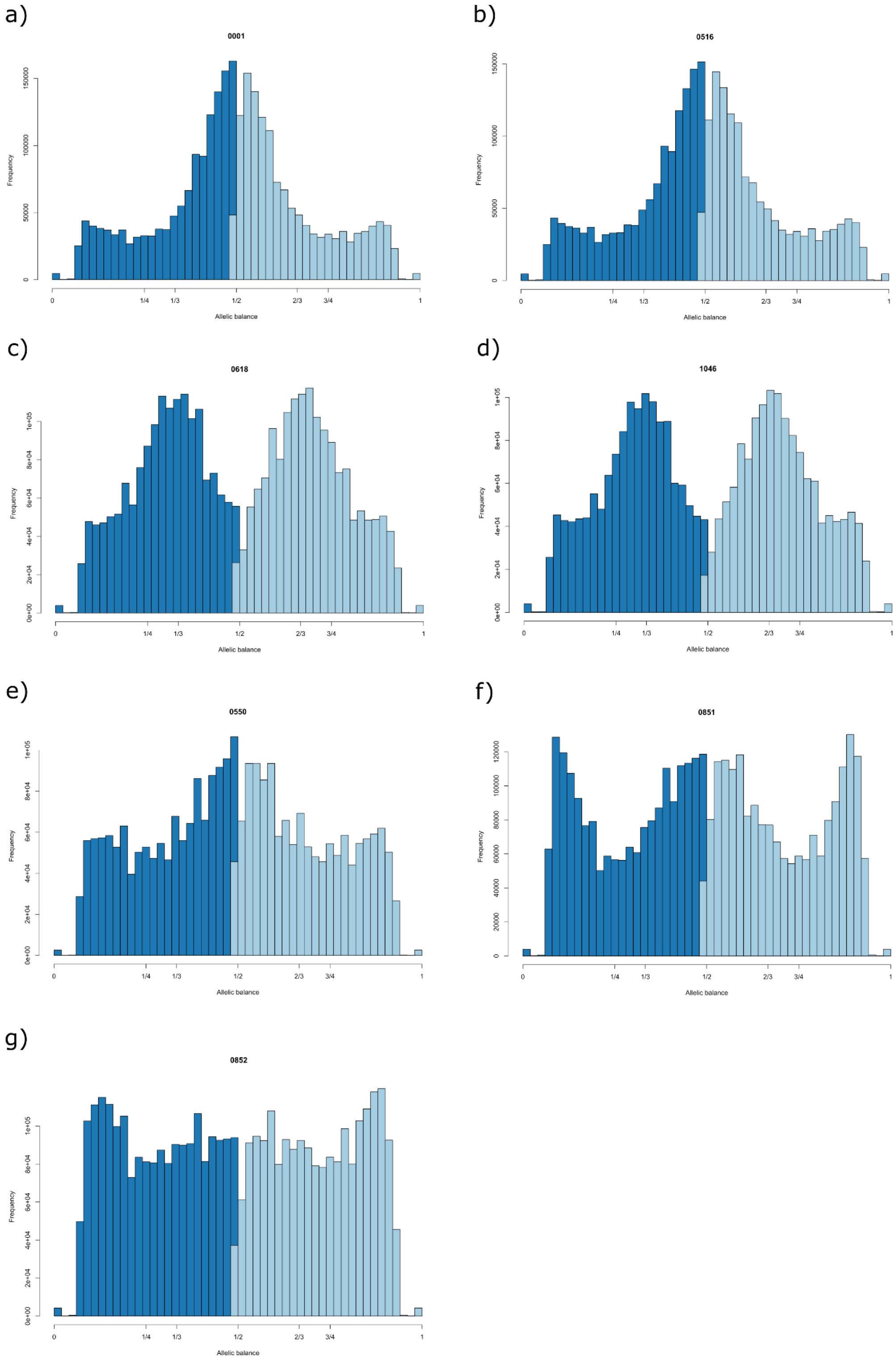


**Figure S1- Allele balance histograms of a 30% subset of SNPs from each *Alnus glutinos*a individual (2n).** Heterozygous SNPs were extracted from the VCF file of a random 30% subset of SNPs for each individual. The fraction of reads supporting each allele (allelic balance) was calculated per site and plotted as a histogram. Panels **a** and **b** show the expected balance for diploids, with a peak at 0.5 indicating equal representation of both alleles. Panels **c** and **d** show histograms characteristic of triploids, with peaks at ~0.3 and ~0.6 indicating the presence of three alleles. Panels **e-g** show individuals with less clearly defined peaks/distributions suggesting ambiguous ploidy.

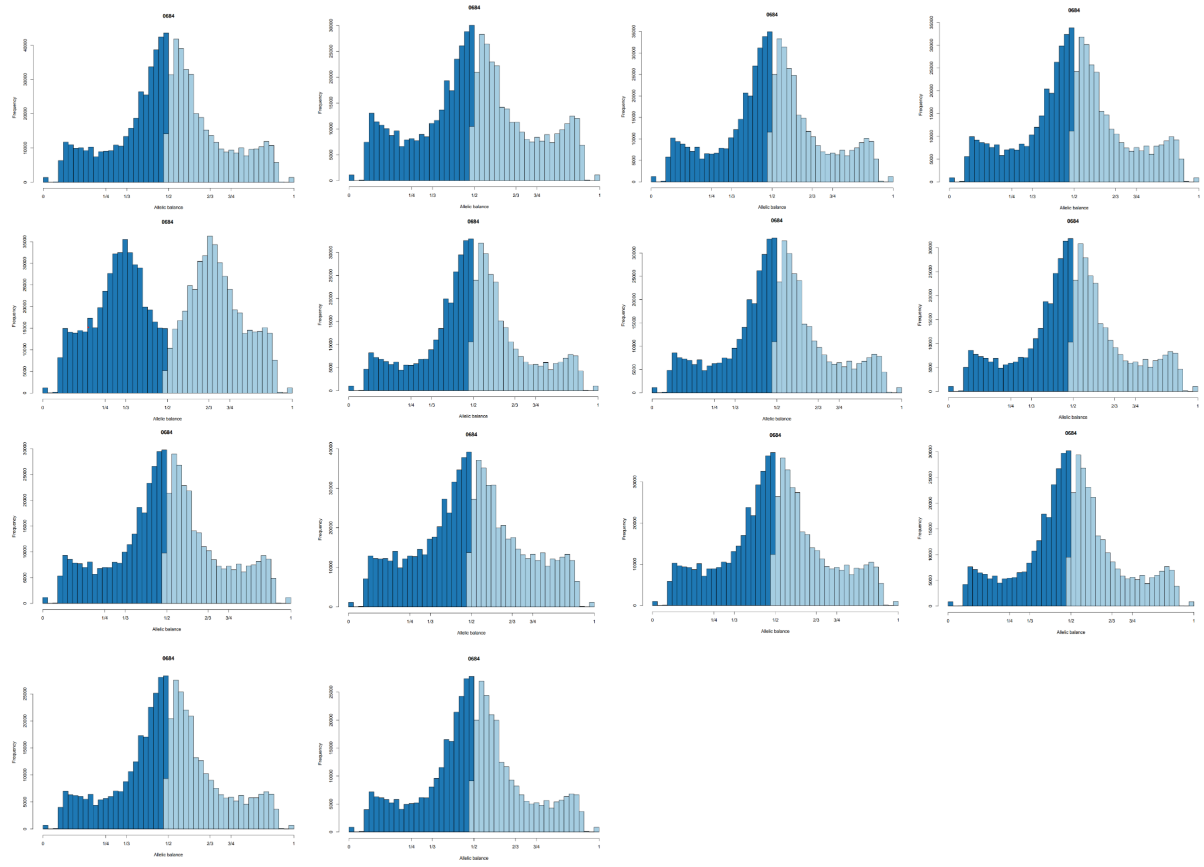


**Figure S2 - Trisomy of individual 0684.** Allele balance histograms consistent with diploidy for all chromosomes except chromosome 5 which shows triploidy. Heterozygous SNPs were extracted from the VCF file for each chromosome in sample 0684, and the fraction of reads supporting each allele (allelic balance) was calculated per site and plotted as a histogram.

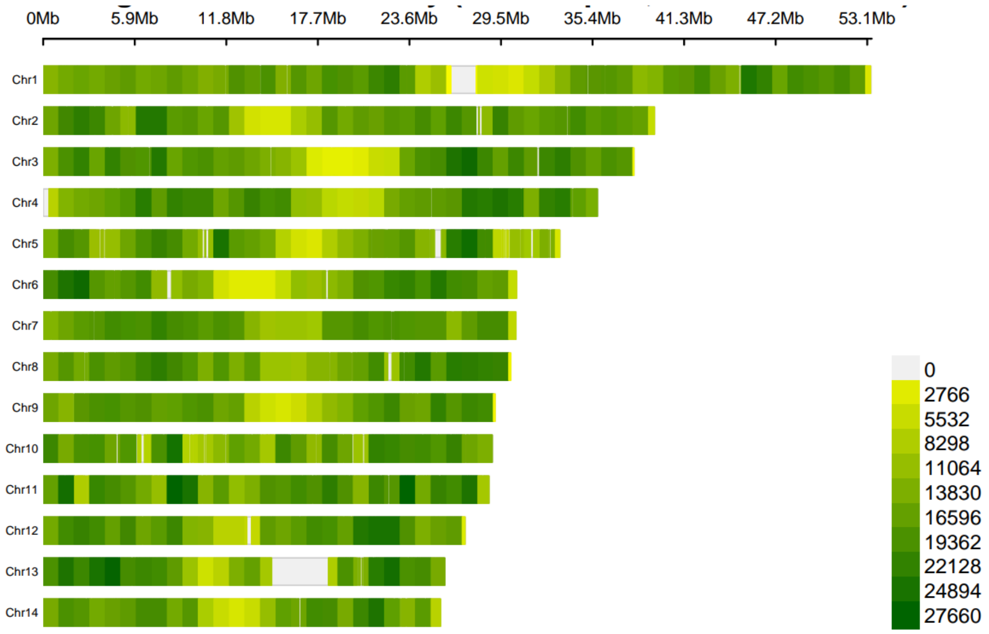


**Figure S3 - SNP density across 14 chromosomes making up the *Alnus glutinosa* genome**. Plot shows ~7.37 SNPs.

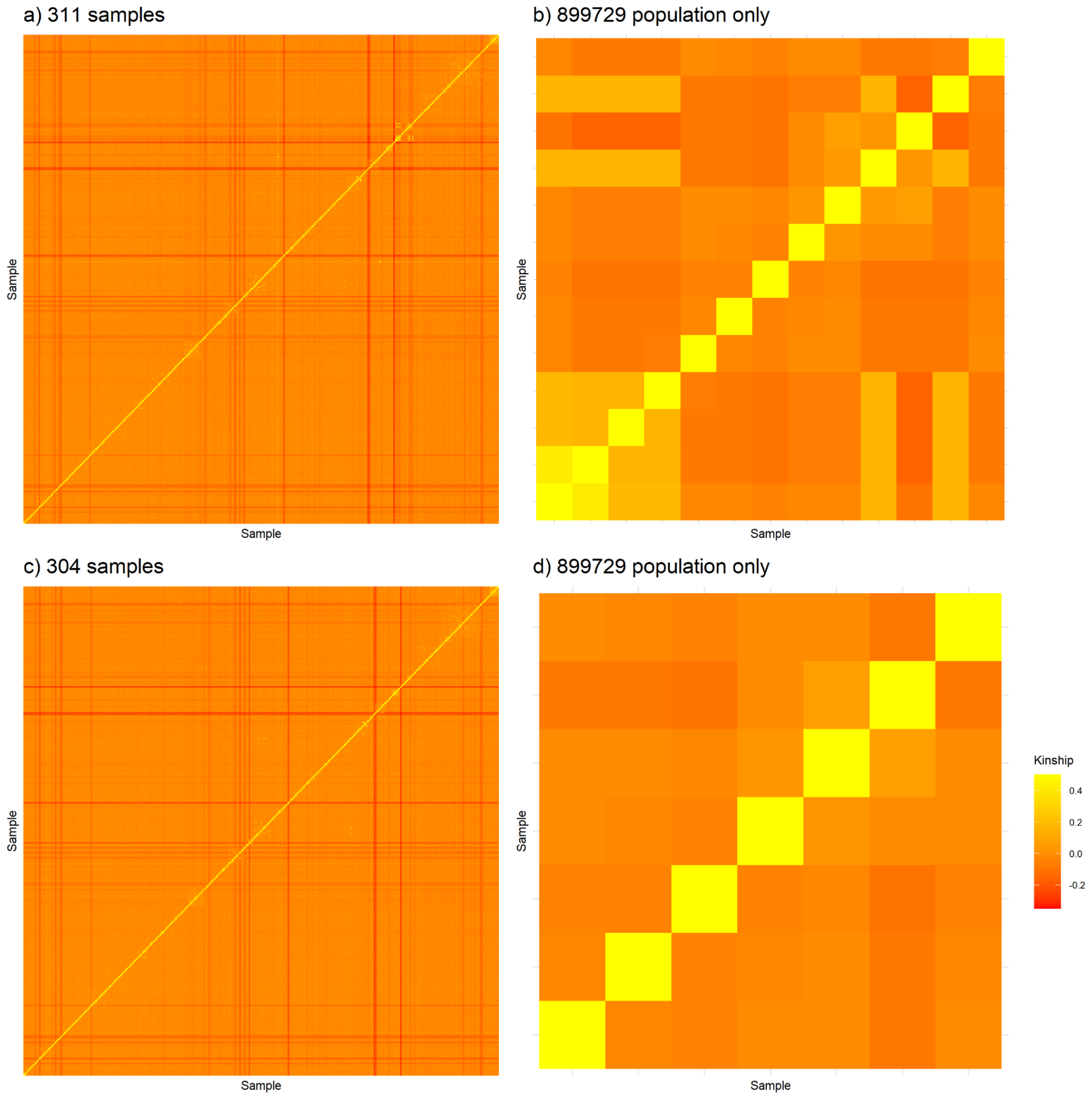


**Figure S4 – Heatmaps of pairwise kinship across 44 UK *Alnus glutinosa* populations.** Polyploids, individuals with ambiguous ploidy, and technical replicates were removed prior to analysis resulting in 311 diploid samples. Plink2 was used to generate a kinship matrix. a shows a heatmap of pairwise kinship across all 311 individuals. b shows intra-population kinship of population 899729, with a subset of six putative diploid hybrids showing high relatedness to each other but low kinship with the remaining samples. c shows a heatmap of pairwise kinship after removal of these six putative hybrid individuals (n=304). d shows intra-population kinship of population 899729 following removal of putative hybrids.

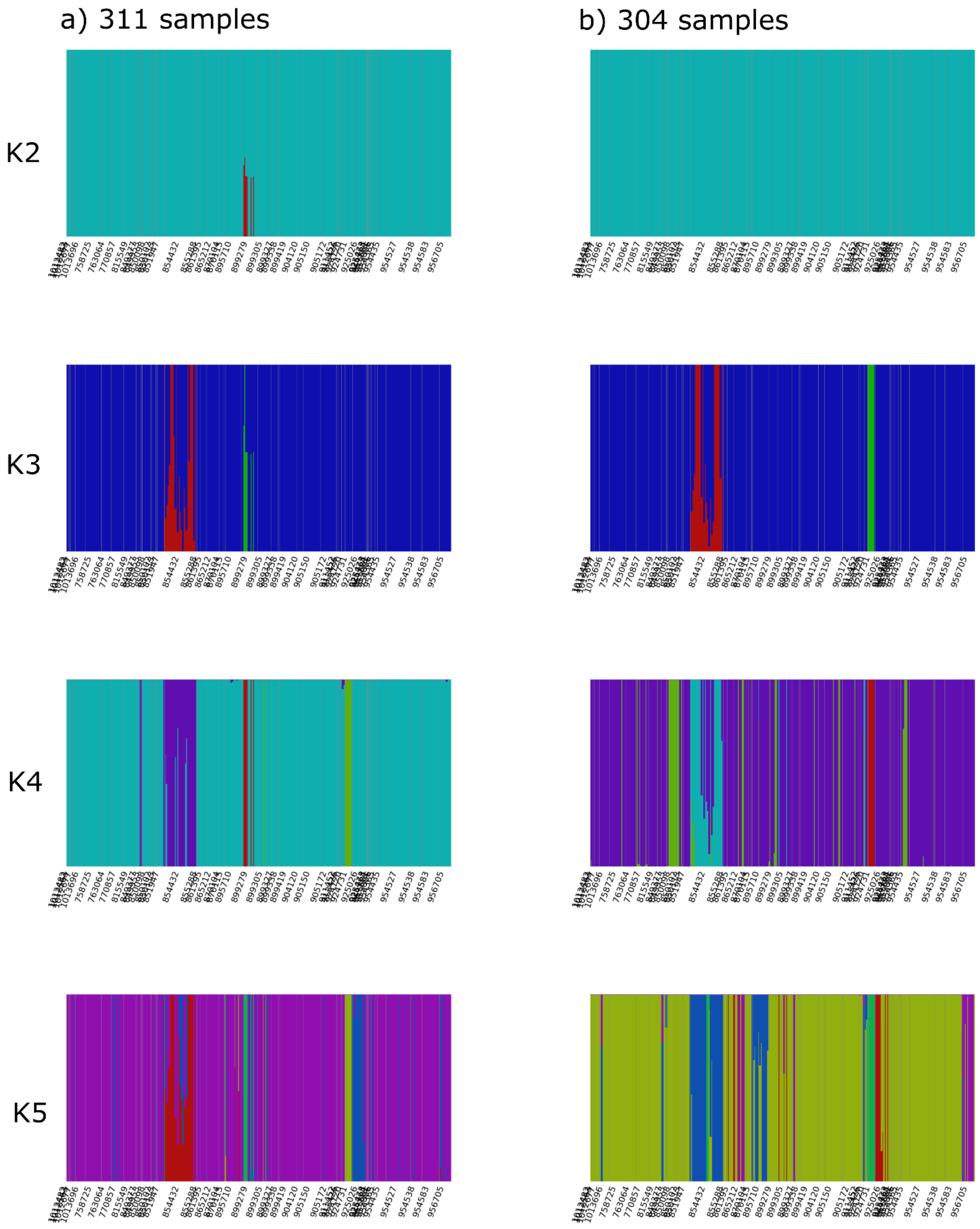


**Figure S5 – FastStructure admixture plots before (a) and after (b) removal of six putative hybrid *Alnus glutinosa* individuals.** Bayesian population structure was inferred using FastStructure from K1-K20 on ~1.5million linkage-filtered SNPs from datasets of 311 and 304 individuals respectively following removal of six putative hybrid individuals.

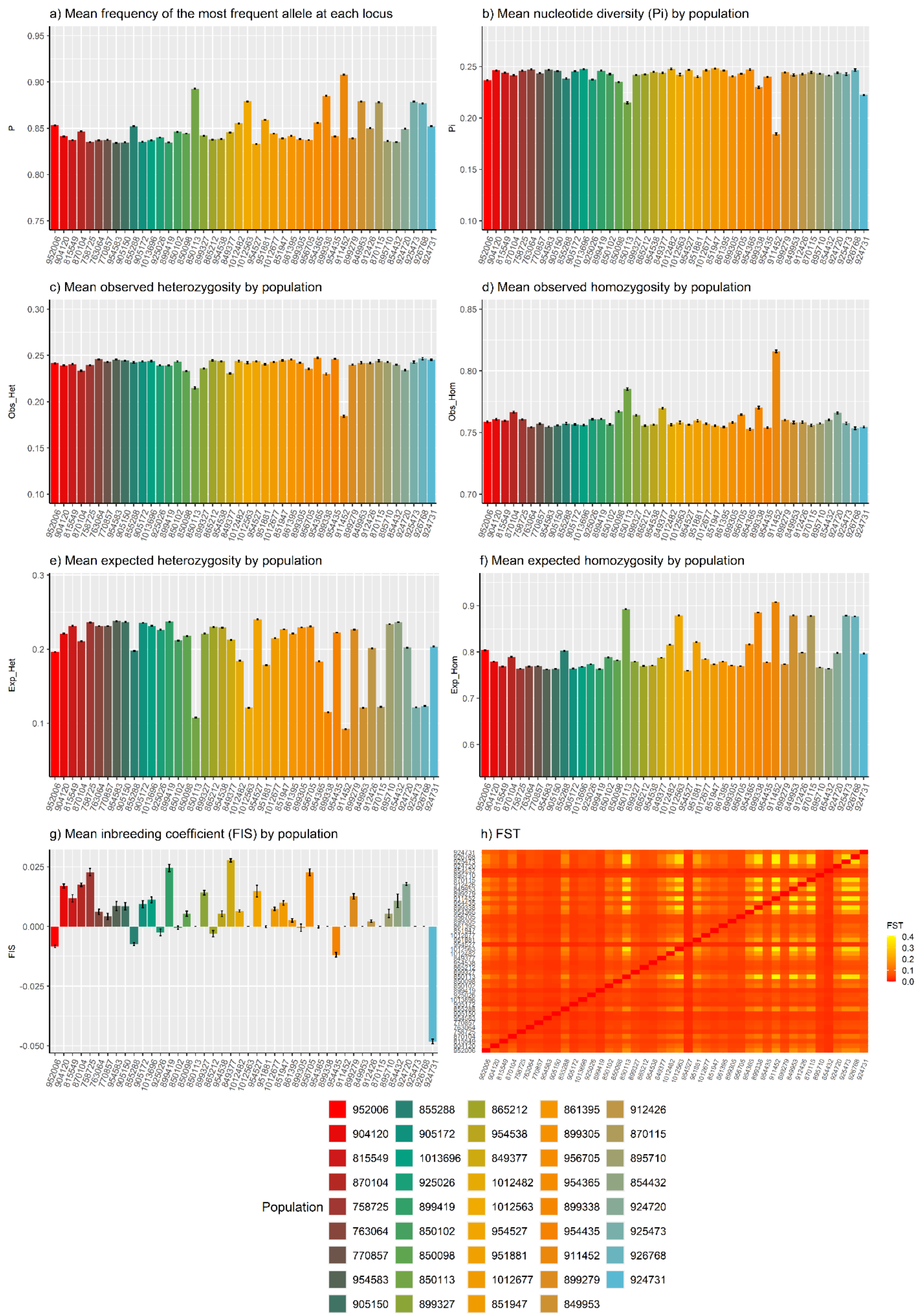


**Figure S6 – Genetic diversity of 44 UK *Alnus glutinosa* populations.** Metrics were calculated using STACKS populations on a 10% random subset of LD-filtered SNPs in 304 individuals across 44 populations. Panels show a) mean frequency of the most frequent allele at each locus (P), b) mean value of Pi (nucleotide diversity, π), c) mean observed heterozygosity (Obs_Het), d) mean observed homozygosity (Obs_Hom), e) mean expected heterozygosity (Exp_Het), f) mean expected homozygosity (Exp_Hom), g) inbreeding coefficient (FIS), h) heatmap of pairwise fixation index ($F_{ST}$) values between each pair of provenances. Populations are ordered by ascending latitude. Note the differing y-axis scales for each plot. π was calculated on biallelic variable sites only, which produces higher values than genome-wide per-base-pair estimates.

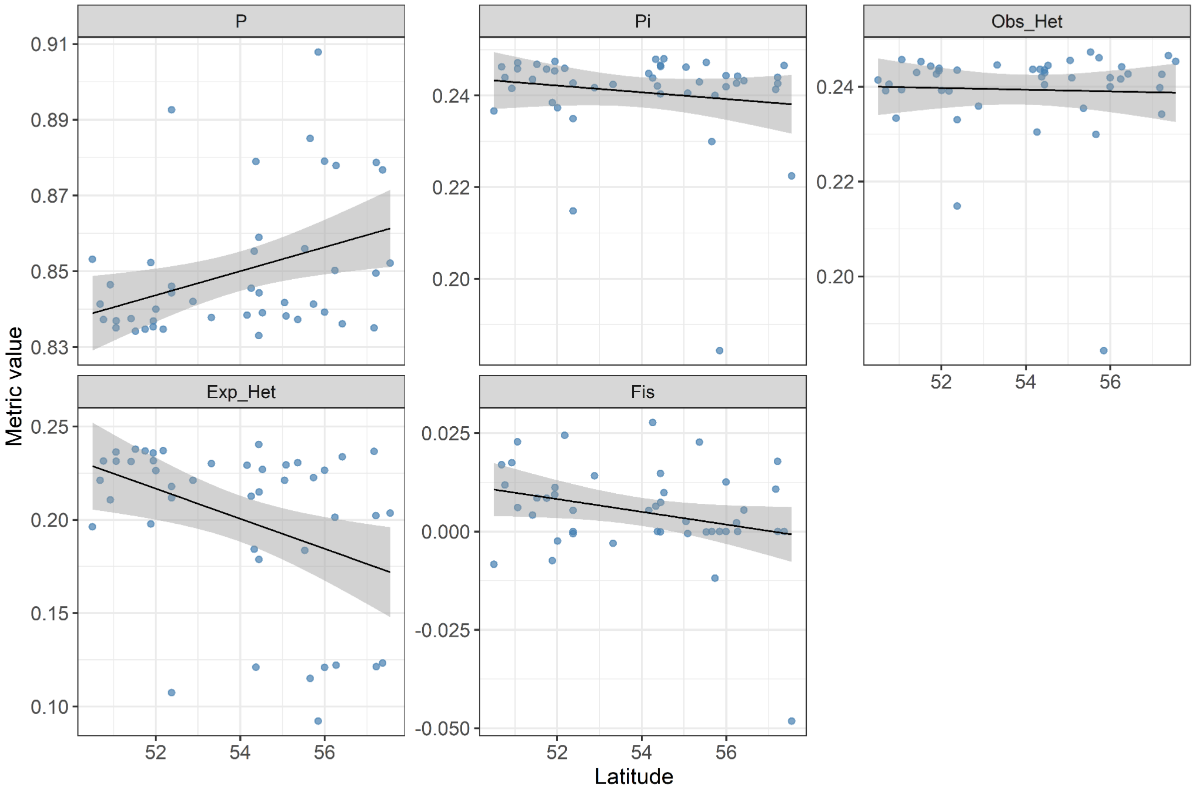


**Figure S7 – Relationship between population latitude and genetic diversity metrics.** Metrics were calculated using STACKS populations on a 10% random subset of LD-filtered SNPs in 304 individuals across 44 populations. Panels show P (mean frequency of the most frequent allele at each locus), Pi (mean value of nucleotide diversity, π), Obs_Het (mean observed heterozygosity), Exp_Het ( mean expected heterozygosity) and Fis (inbreeding coefficient (FIS). The latitude of each population was regressed on each diversity metric, and p-values were estimated using F-tests (Table S2).

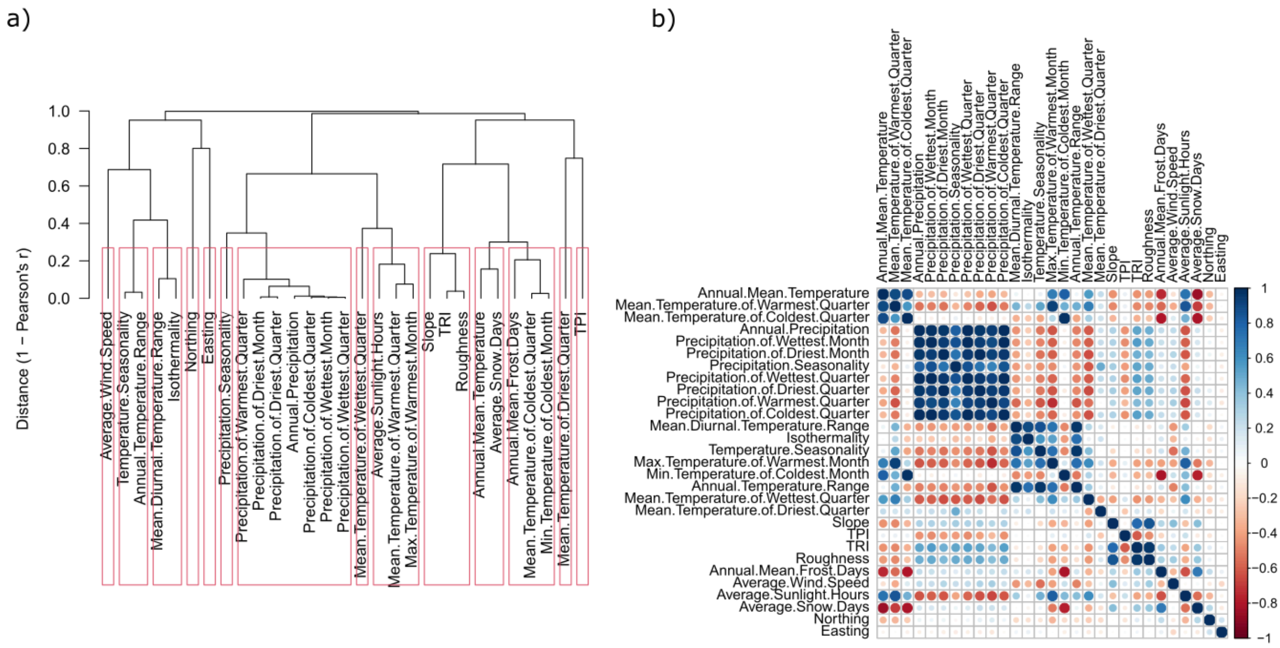


**Figure S8: Collinearity between 29 environmental and terrain variables extracted from the sites of 44 UK *Alnus glutinosa* populations.** Variables included 19 CHELSA climatologies, 4 MetOffice variables, and 6 terrain characteristics. **a,** Plot produced by the R package *virtualspecies,* grouping variables with a correlation coefficient greater than 0.7. **b,** Pairwise correlation coefficients plotted using the R package *corrplot*.

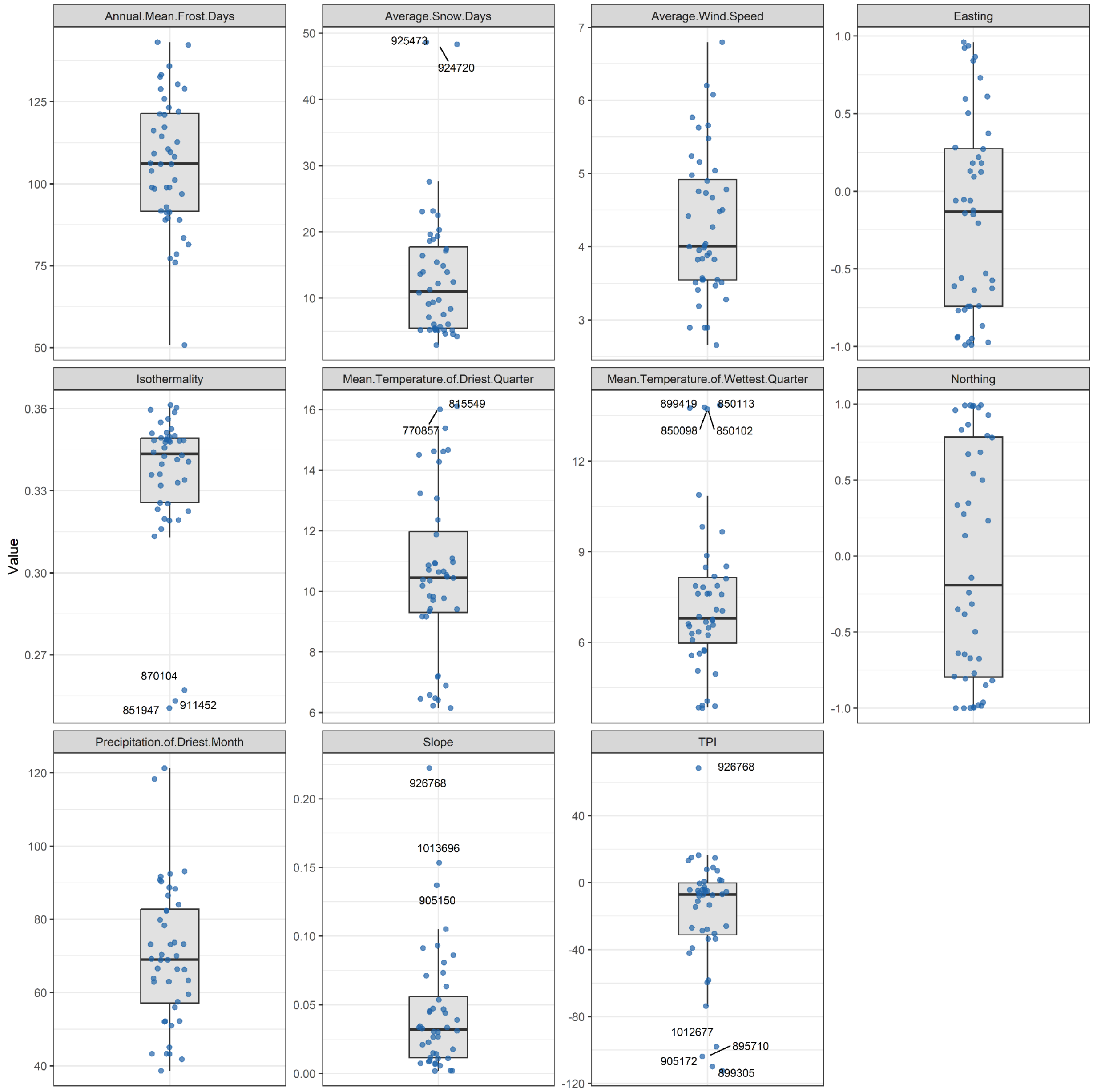


**Figure S9: Distribution of the 11 environmental variables across the 44 UK *Alnus glutinosa* populations used in the GEA.** Each point represents a population's local conditions for each variable, with populations showing outlying values labelled (lowest and highest quartiles).

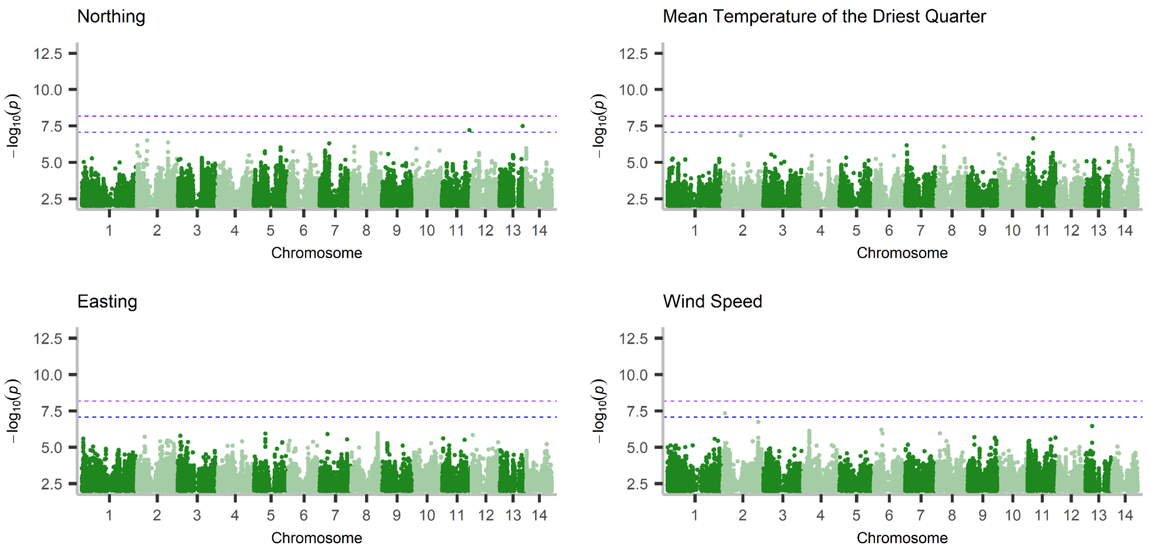


**Figure S10– Manhattan plots following GEA analysis on 304 UK *Alnus glutinosa* individuals across 44 populations**. Models were performed on 7.38 million SNPs at K2. Horizontal lines indicate modified-Bonferroni (blue) and Bonferroni (purple) significance thresholds. Variables shown are those without significance according to the FDR threshold. Plots for variables with significance are available in Figure 3 in the main text.

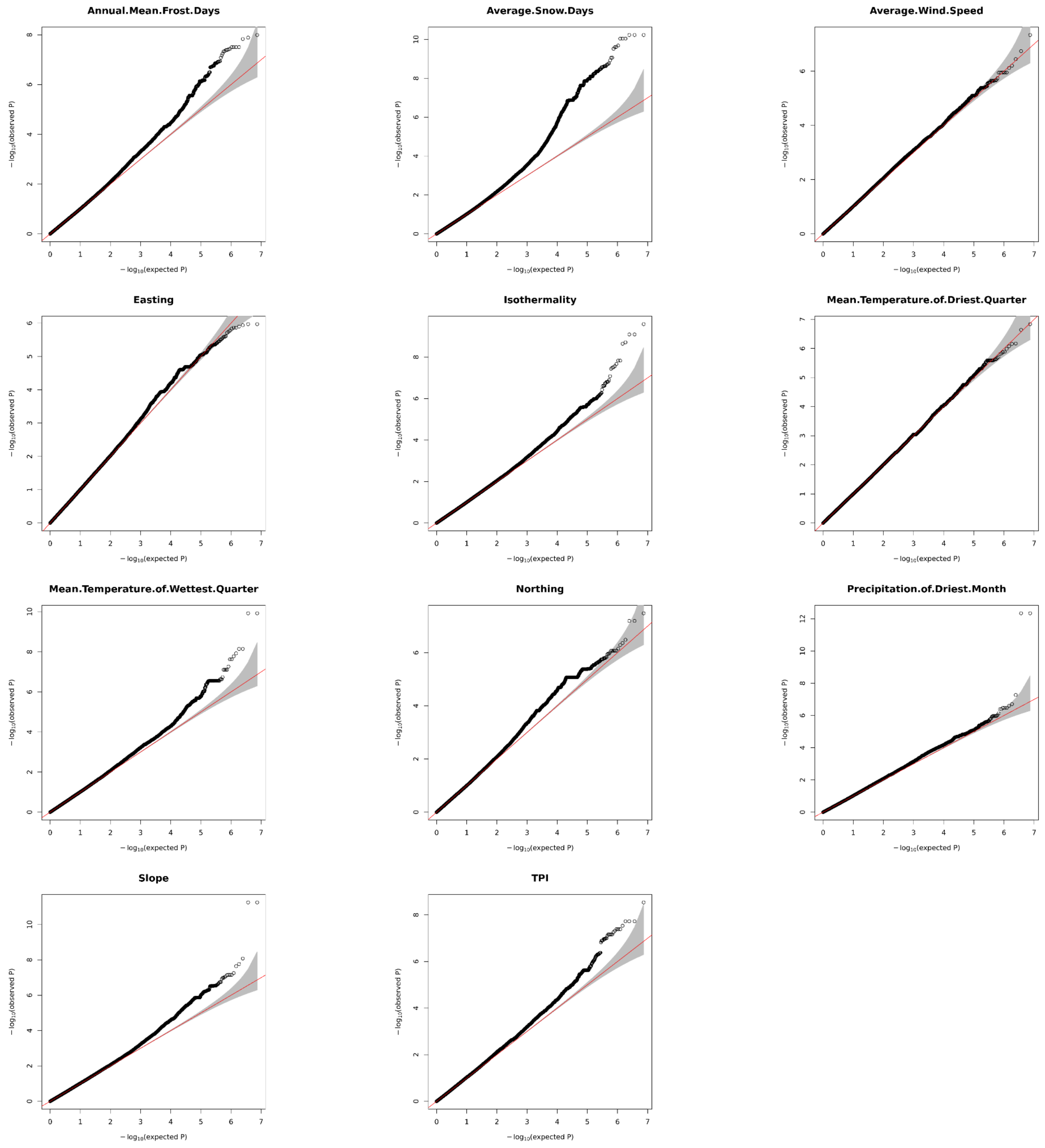


**Figure S11– QQ-plots following LFMM analysis at K2 on 304 UK *Alnus glutinosa* individuals from 44 populations**. LFMM were performed on 7.38 million SNPs and 11 environmental variables.

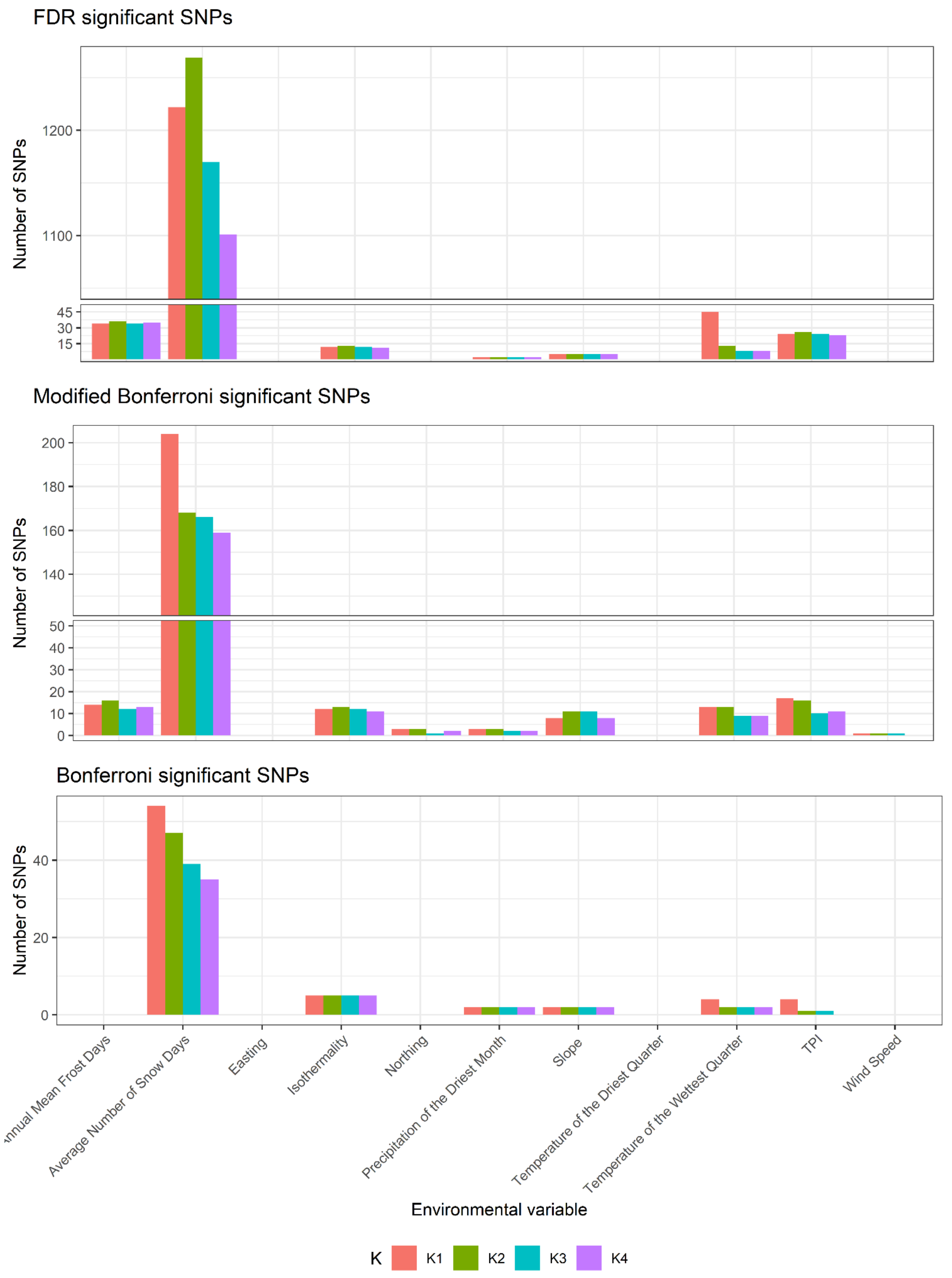


**Figure S12 – Number of significant SNPs associated with each environmental variable following LFMM analysis at K1-K4.** LFMMs were performed on 7.38 million SNPs to 11 environmental variables at K1 to K4. Significant SNPs according to a false discovery rate of 5% were counted along with significance according to Bonferroni and Modified Bonferroni thresholds (dividing by the total number of SNPs and haplotype blocks respectively).

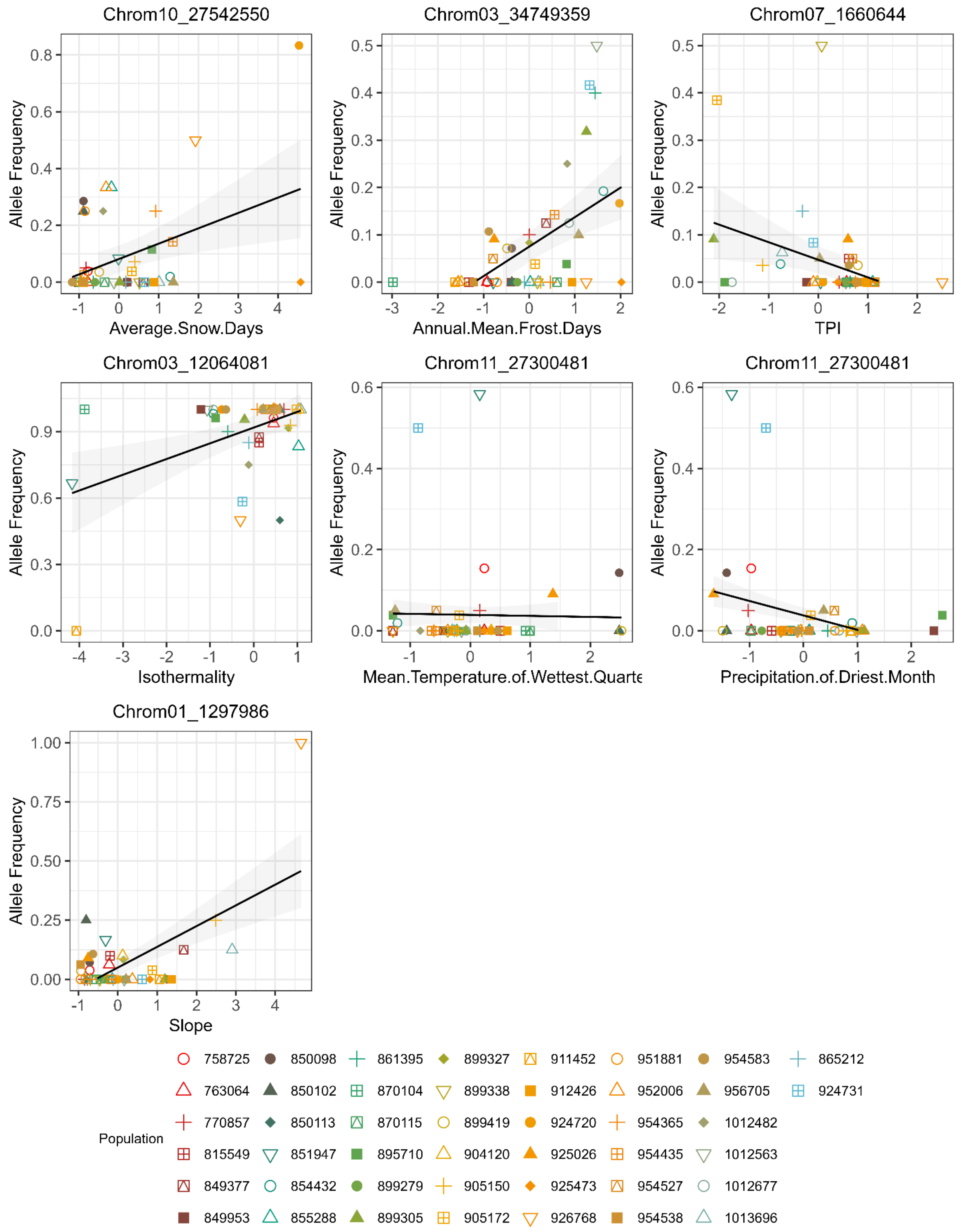


**Figure S13 – Population-level allele frequencies of the SNPs most significantly associated with the environment at each source location.** Each panel shows the allele frequency of the single SNP with the lowest p-value among all significant SNPs associated with that environmental variable. Allele frequencies were calculated using PLINK.

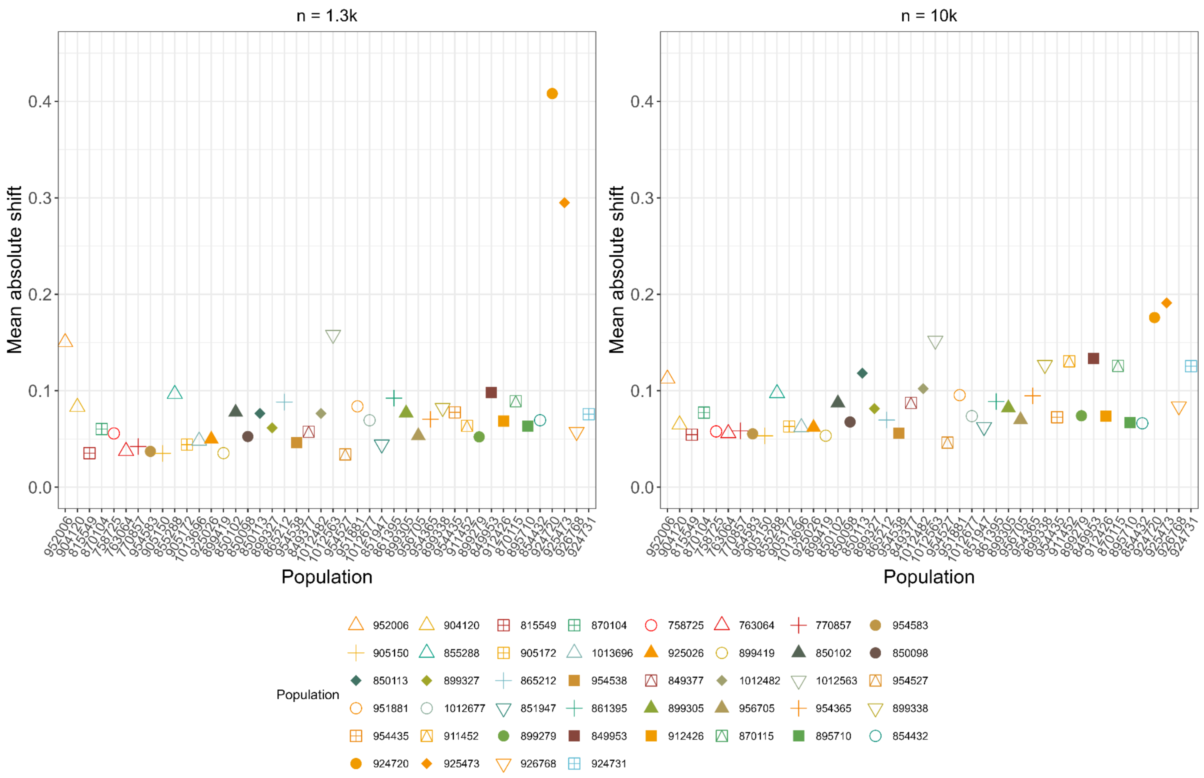


**Figure S14 – Predicted present day maladaptation of 44 UK *Alnus glutinosa* populations to local environmental conditions.** Genetic change was estimated in the *AlleleShift* R package to determine the mean allele frequency change (allele shift) required for SNPs to be present at the optimum frequency per population in their own source, local conditions. Analysis was performed on **a,** the 1,364 FDR-significant climate-associated SNPs and **b,** the 10,000 SNPs with the lowest p-values following LFMM2 analysis. Populations are ordered by latitude. Latitude was a significant predictor of present-day allele shift for both SNP sets (AlleleShift1K ~ Latitude: $F_{1,42} = 6.15$, $p = 0.017$, $R^2 = 0.13$; AlleleShift10K ~Latitude: $F_{1,42} = 13.75$, $p = <0.01$, $R^2 = 0.25$). For estimations using 1K SNPs, the relationship between allele shift and latitude was no longer significant following the exclusion of populations 924720 and 925473 ($F_{1,40} = 1.03$, $p = 0.32$, $R^2 = 0.02$).

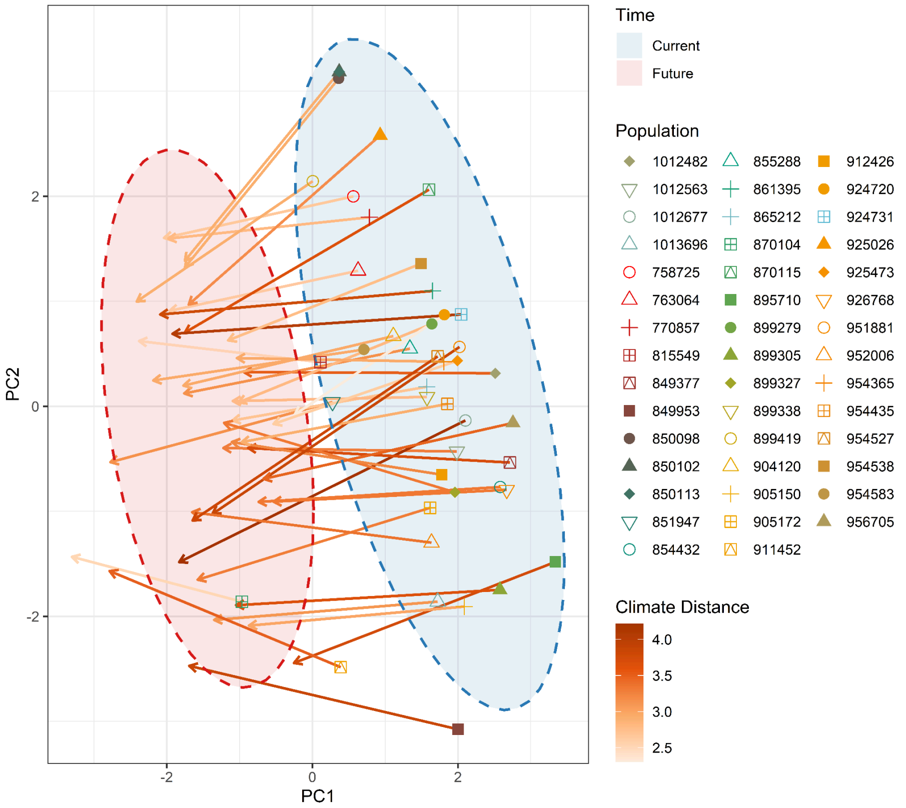


**Figure S15 – Climatic difference between current and future environmental conditions in a worst-case climate change scenario at the site of each of the 44 UK *Alnus glutinosa* populations.** PCA was performed on ten uncorrelated environmental variables as included in the genomic offset and AlleleShift estimations. Ellipses are coloured by time period, with blue for current and red for future. Points are coloured and shaped by population. Arrows connect populations across time points, and are coloured according to the climate distance calculated from PC1-PC5, with deeper colours representing greater environmental change.

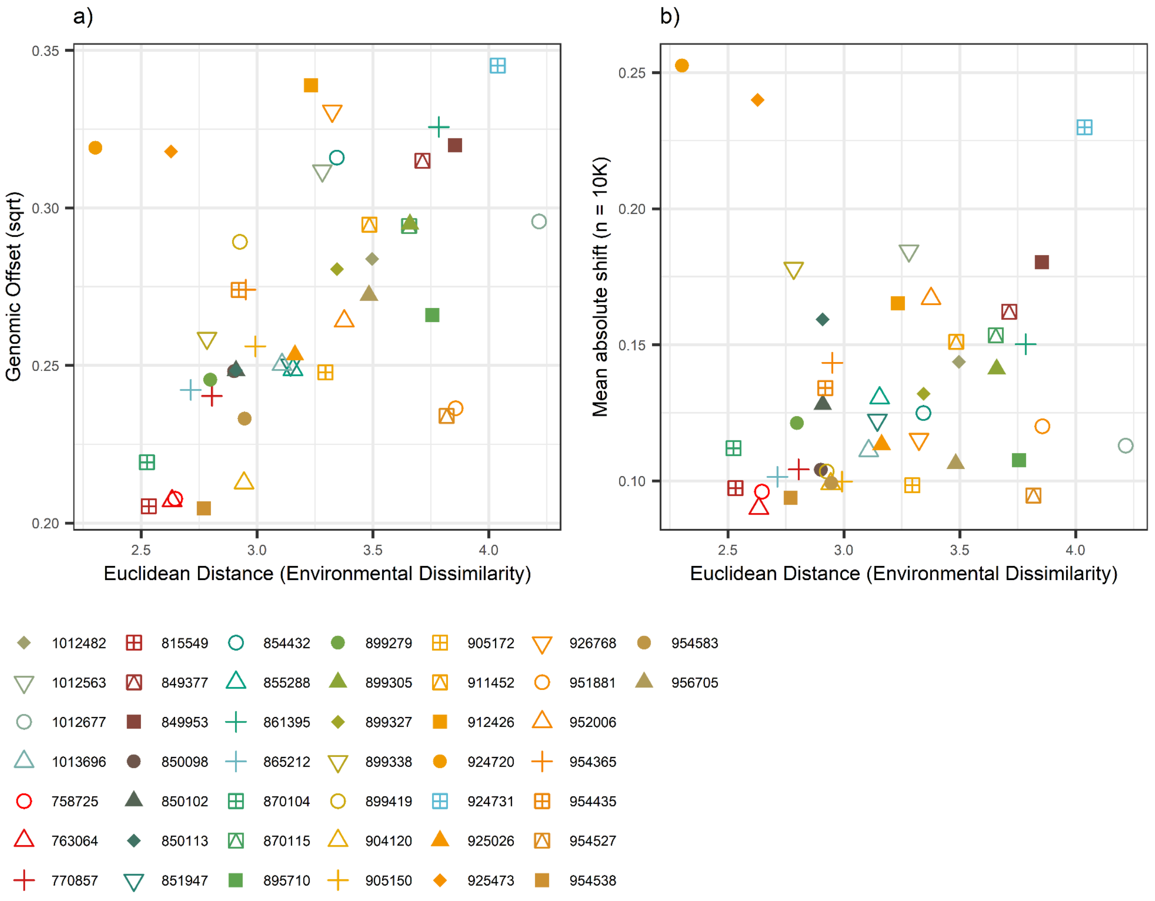


**Figure S16 – Predicted environmental and genetic change of 44 UK *Alnus glutinosa* populations in a worst-case climate change scenario at each seed source location.** Differences between current and future predicted climates were calculated as the Euclidean distance in PCA space from PC one to PC five. **a,** Environmental dissimilarity per population against genomic offset of each population estimated in the *LEA* package on 1.49 million LD-filtered SNPs. **b,** Environmental dissimilarity per population against the mean allele frequency change (allele shift) required for 10,000 SNPs with the lowest p-value following LFMM2 to be at an optimum frequency per population. Euclidean distance was a significant predictor of genomic offset (GenomicOffset ~ ClimateDissimilarity: $F_{1,42} = 12.91$, $p < 0.01$, $R^2 = 0.24$), but not allele shift (10K SNPs) (AlleleShift10K ~ ClimateDissimilarity: $F_{1,42} = 0.33$, $p = 0.57$, $R^2 = 0.01$).

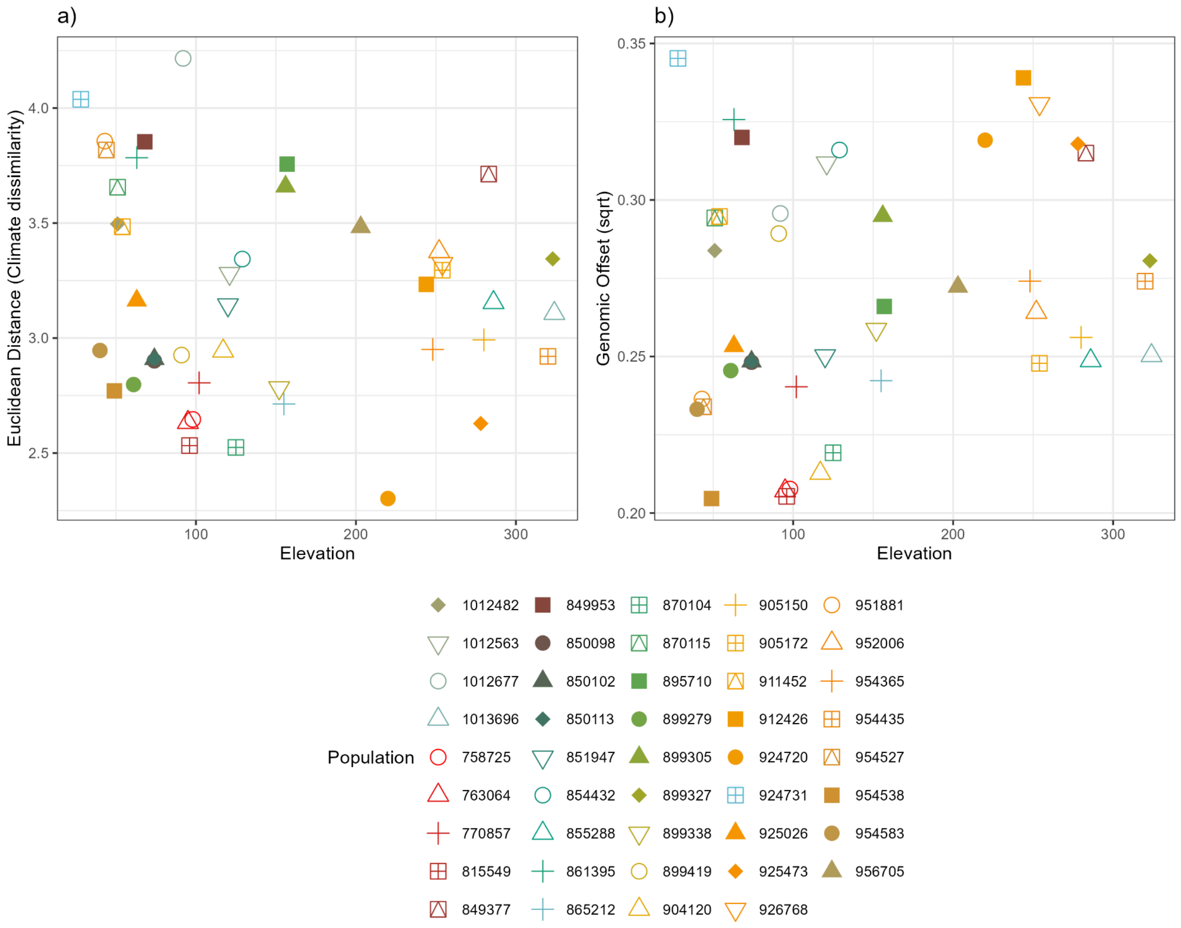


**Figure S17 – Predicted environmental and genetic change of 44 UK *Alnus glutinosa* populations in a worst-case climate change scenario against elevation at each seed source location. a,** Environmental dissimilarity between current and future local conditions per population against elevation of each population. Differences between current and future predicted climates were calculated as the Euclidean distance in PCA space from PC one to PC five. **b,** relationship between genetic offset and elevation for each population. The genetic.offset() command within the *LEA* package was used to estimate the genetic offset of 1.49 million LD-filtered SNPs from 304 individuals across 44 populations. Elevation was not a significant predictor of either metric (EuclideanDistance ~ elevation: $F_{1,42} = 0.88$, p =0.35, $R^2 = 0.02$; GenomicOffset ~ elevation:($F_{1,42} = 1.96$, p =0.169, $R^2 = 0.045$)

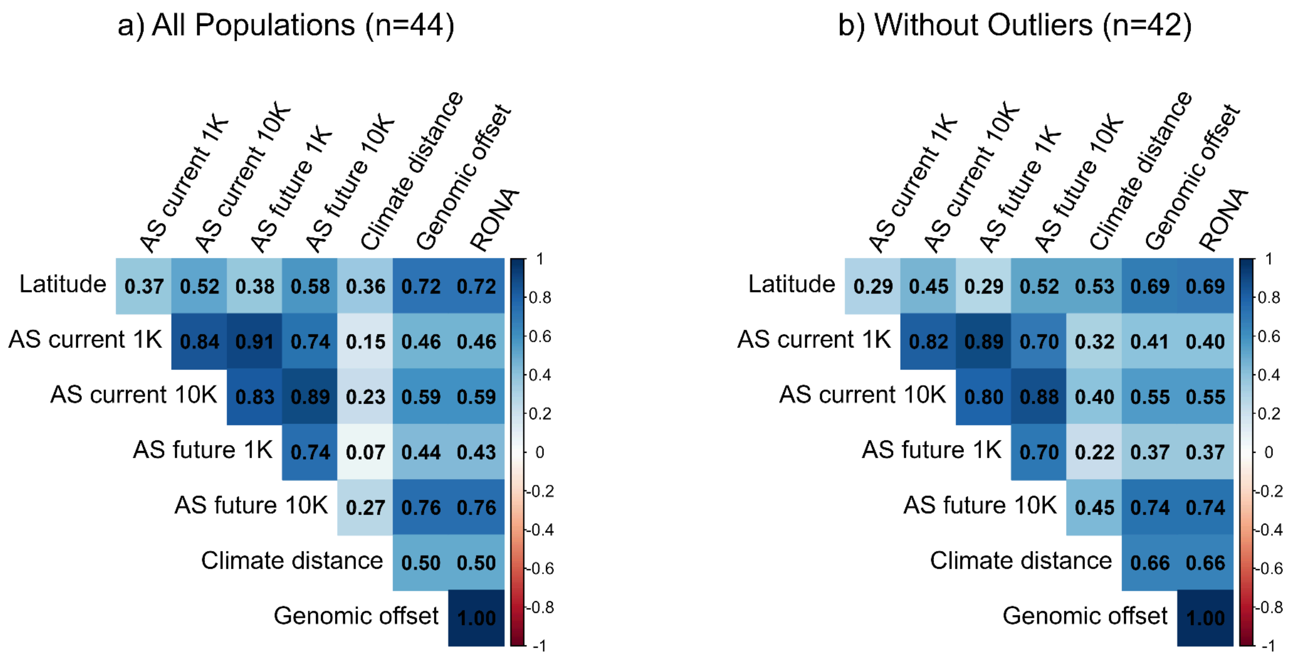


**Figure S18 - Correlation between estimates of genetic maladaptation, environmental dissimilarity, and latitude**. **a,** correlations when all populations were included. **b,** correlations when two outlying populations (924720 and 925473) were removed. Metrics include: latitude; AS current 1K = allele shifts to present-day local climates calculated using SNPs above an FDR significance threshold in GEA; AS current 10K = allele shifts to present-day local climates calculated using the top 10,000 SNPs by p-value following GEA; AS future 1K = allele shifts to predicted future climates calculated using SNPs above an FDR significance threshold in GEA; AS future 10K = allele shifts to predicted future climates calculated using the top 10,000 SNPs by p-value following GEA; Climate distance = Euclidean distance in PCA space between present-day and future predicted local environmental conditions per population; Genomic offset = genetic gap between present day and future climates calculated with 1.49million genome-wide SNPs; RONA = modified risk of non-adaptedness calculated with 1.49million genome-wide SNPs. For details of how these estimates were calculated, please see Methods section.

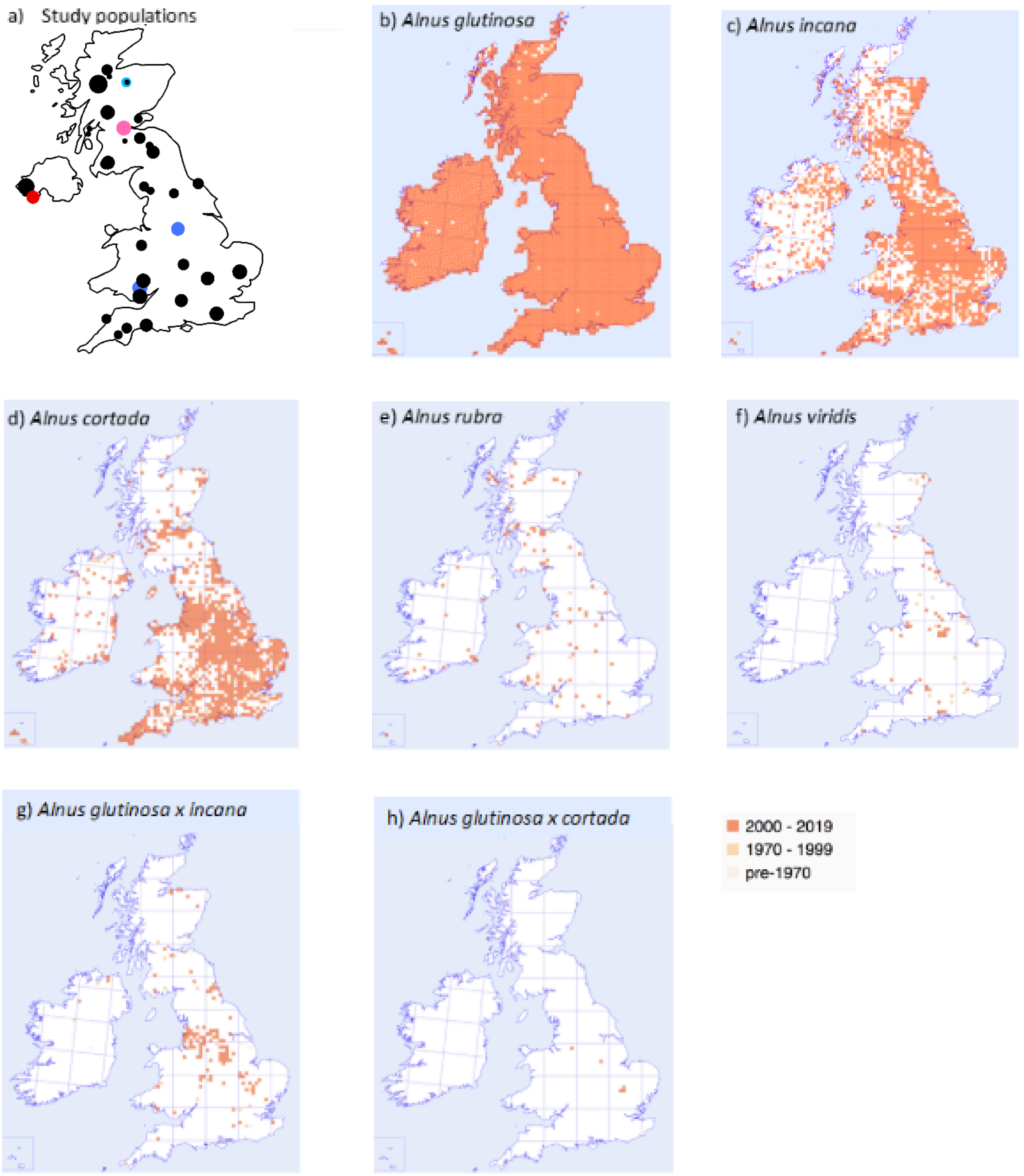


**Figure S19 - Maps from the Botanical Society of Britain and Ireland showing the distribution of UK *Alnus* species** (Botanical Society of Britain & Ireland 2026)**. a)** shows the location of the 44 populations used in this study with dark blue for populations with triploids, light plink for individual 0550 (ambiguous ploidy likely caused by high rates of missingness), red for populations containing individuals for which ploidy could not be determined, dark pink for diploid hybrids, and light blue for the population with a trisomic individual. Maps **b-h** show *Alnus* species distribution coloured according to recording date, with **b)** for *A.glutinosa*, **c)** for *A.incana*, **d)** for *A.cortada*, **e)** for *A.rubra*, **f)** for *A.viridis*, **g)** for *A.glutinosa x A.incan*a hybrids, and **h)** for *A.glutinosa x A.cortada* hybrids.
